# Mapping calcium dynamics in a developing tubular structure

**DOI:** 10.1101/2020.10.16.342535

**Authors:** Jorgen Hoyer, Morsal Saba, Daniel Dondorp, Kushal Kolar, Riccardo Esposito, Marios Chatzigeorgiou

## Abstract

Calcium is a ubiquitous and versatile second messenger that plays a central role in the development and function of a wide range of cell types, tissues and organs. Despite significant recent progress in the understanding of calcium (Ca^2+^) signalling in organs such as the developing and adult brain, we have relatively little knowledge of the contribution of Ca^2+^ to the development of tubes, structures widely present in multicellular organisms. Here we image Ca^2+^ dynamics in the developing notochord of *Ciona intestinalis*. We show that notochord cells exhibit distinct Ca^2+^ dynamics during specific morphogenetic events such as cell intercalation, cell elongation and tubulogenesis. We used an optogenetically controlled Ca^2+^ actuator to show that sequestration of Ca^2+^ results in defective notochord cell intercalation, and pharmacological inhibition to reveal that stretch-activated ion channels (SACs), inositol triphosphate receptor (IP3R) signalling, Store Operated Calcium Entry (SOCE), Sarco/endoplasmic reticulum Ca^2+^-ATPase (SERCA) and gap junctions are required for regulating notochord Ca^2+^ activity during tubulogenesis. Cytoskeletal rearrangements drive the cell shape changes that accompany tubulogenesis. In line with this, we show that Ca^2+^ signalling modulates reorganization of the cytoskeletal network across the morphogenetic events leading up to and during tubulogenesis of the notochord. We additionally demonstrate that perturbation of the actin cytoskeleton drastically remodels Ca^2+^ dynamics, suggesting a feedback mechanism between actin dynamics and Ca^2+^ signalling during notochord development. This work provides a framework to quantitatively define how Ca^2+^ signalling regulates tubulogenesis using the notochord as model organ, a defining structure of all chordates.

## Introduction

Tubular structures are essential in biological organisms, playing a critical role in the structural support of surrounding tissues, as well as the transport and exchange of fluids and molecules^1^. Tube morphogenesis is a highly complex process that is orchestrated by a series of molecular and cellular changes^2–6^. These cellular changes include cell shape alteration, cell proliferation, mesenchymal-epithelial transition (MET), cell polarization, cell migration and cell differentiation^3, 6^. Coordinating collective changes in cell state and cell behaviour requires spatially and temporally versatile yet robust signalling. Calcium (Ca^2+^) signalling is one of the primary mechanisms employed^7–11^. At the molecular level, it regulates the expression of large batteries of genes using Ca^2+^ responsive transcriptional cascades^12–14^. At the cellular level, processes regulated by Ca^2+^ signalling include cell proliferation^15, 16^, differentiation^17^, migration^18, 19^ and morphogenesis^20^. The importance of Ca^2+^ signalling in embryonic development has been recognized for over 20 years^19, 21–24^, motivating Ca^2+^ imaging studies in a range of organisms including *C. elegans*^25^, Drosophila^26–29^, the tunicates *C. robusta^30^’*^33^ and *Oikopleura dioica*^34^, zebrafish^35–39^, xenopus^18, 40–44^, butterfly^45^ and mouse^46^. However, an important gap in our current knowledge is how Ca^2+^ signalling dynamics mediate the morphogenetic processes that give rise to tubular structures such as blood vessels, renal tubules, the neural tube and the notochord.

The notochord plays an essential role as a signalling centre in the organization of the embryonic body plan of chordates. During early embryonic development in vertebrates, the notochord is required for patterning of the neural tube, formation of the paraxial mesoderm, establishment and maintenance of left-right asymmetry^47^ and formation of key organs such as the heart and blood vessels^48^. Abnormal notochord development therefore leads to global embryonic malformations^49–54^. Furthermore, an aggressive type of tumour, chordoma, develops from activated notochord cells ^55–57^.

*Ciona intestinalis* is an emerging model to study the evolution, development and function of the notochord in chordates. At the molecular level, gene regulatory networks that are required for the development and function of the notochord have been identified^58–66^. Comparative studies between tunicate species have increased our understanding of the evolutionary changes that took place in the notochord at the genetic and cellular levels^67–70^. Finally, multiple studies have shed light on the cell biological basis of notochord morphogenesis^69, 71–81^.

*C. intestinalis* embryogenesis is divided into 26 stages, starting from the zygote and ending with the hatching larva^82^. Concurrent with gastrulation (stages 10-13), the notochord cells divide twice to result in 40 cells that no longer divide, are geometrically identical, and arranged in a flat sheet. The subsequent steps of notochord development involve a series of morphogenetic processes including cell shape changes, lumen formation and tissue reconfiguration (Figure 1A): during the neurula period, the sheet of notochord cells invaginates to form a rod. Between late neurula (stage 16) and early tailbud stages (stage 20), notochord cells intercalate radially and medio-laterally via convergent extension^69, 80^. As a result, the mid-tailbud embryos (stage 21) have a columnar notochord composed of 40 cells aligned in a single file, resembling a stack of coins. Subsequent tubulogenesis is initiated. This process begins with cell elongation along the antero-posterior axis and polarization (Stage22-23), followed by appearance of apical domains and extracellular lumen pockets that emerge between neighbouring cylindrical cells (stage 24). Finally, during stage 25 and stage 26, notochord cells perform bidirectional crawling movements to allow the neighbouring lumen pockets to fuse together, thus enabling the cells to adopt an endothelial-like morphology.

**Figure 1:**
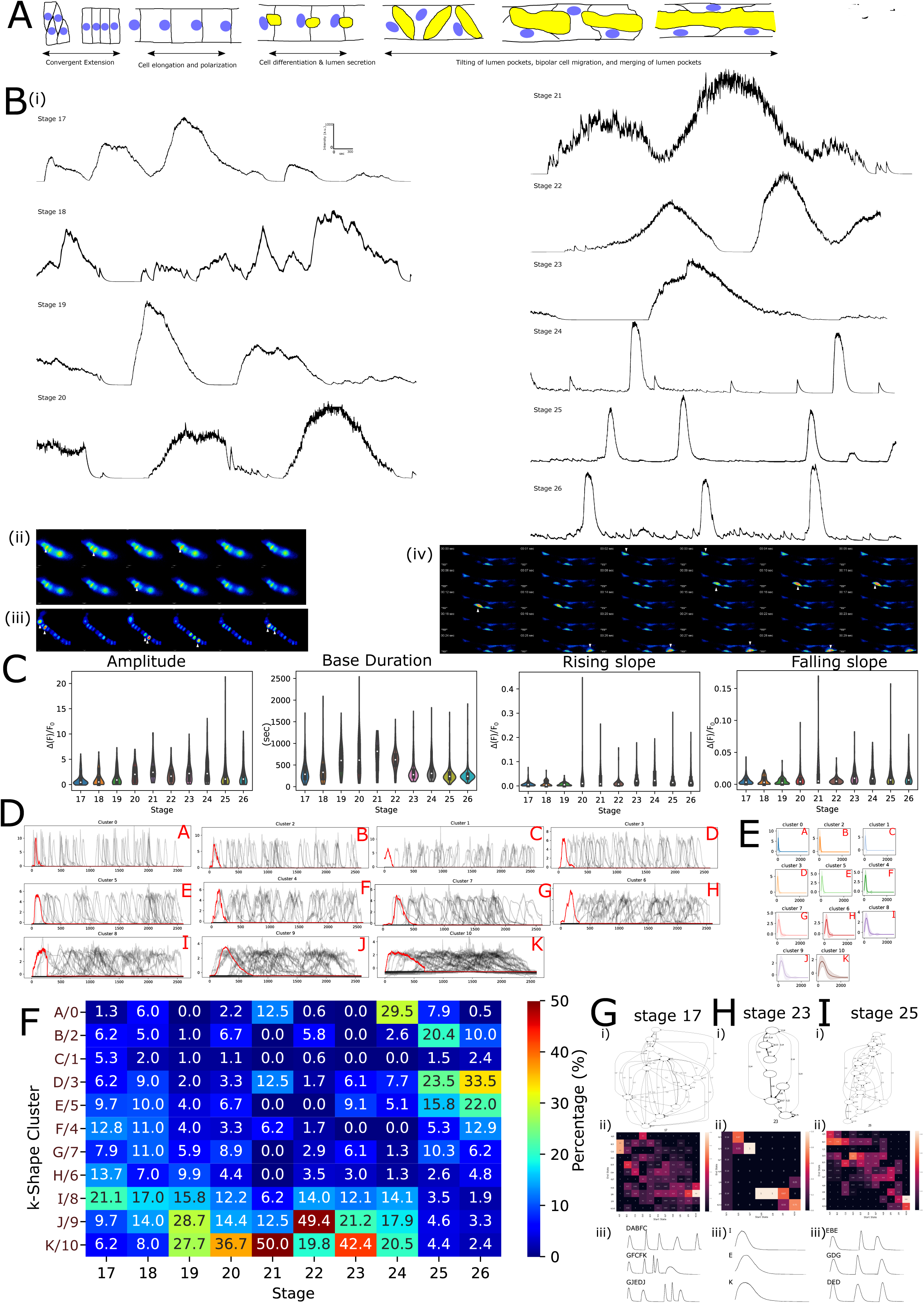
The developing notochord exhibits dynamically changing Ca^2+^ activity profiles. A) Illustration of the key morphogenetic steps during notochord formation. B) i) Example traces of Ca^2+^ activity at different developmental stages. Montage of pseudocoloured frames from ii) Stage 17 embryo expressing GCaMP6s under the control of Brachyury promoter showing Ca^2+^ activity. iii) Stage 24 tailbud embryo showing Ca^2+^ activity in a number of cells. iv) Late stage 25 embryo showing Ca^2+^ activity in notochord cells exhibiting bipolar cell migration. Cells with increasing activity shown with white arrowhead. C) Peak features of wild-type embryos imaged at different developmental stages (17-26). 4-14 embryos were used per developmental stage. (see Sup. Table1) D) k-Shape clusters with centre of cluster transient shown in red and randomly sampled traces from each cluster shown in black. E) Mean traces of all individual traces contained in each cluster shown with S.E.M. F) k-Shape cluster composition (unit is %) colour coded with blue to red (low to high percentages). G) Stage 17, H) Stage 23, I) Stage 25: i) Markov-Chain graphs from different developmental stages. ii) Markov-Chain matrixes reveal the difference in the complexity of the transitions taking places at different stages. ii) Example traces generated from the Markov Chain model illustrating differences in activity patterns during development.

The main goals of our study are to: 1. determine the extent to which Ca^2+^ dynamics are present throughout development in the notochord 2. quantitatively define the relationship between the complexity of Ca^2+^ dynamics and specific morphogenetic events occurring in the developing notochord 3. directly perturb Ca^2+^ signalling, via optogenetics and pharmacological means, to test its requirements in convergent extension and three-dimensional (3D) reorganization of the cytoskeleton during notochord development.

For data analysis, we have applied novel approaches to dissect the complex spatiotemporal Ca^2+^ dynamics of a developing tubular structure, the notochord of C. intestinalis. Our experimental data and analysis revealed that Ca^2+^ activity is present throughout notochord development, from cell intercalation until the formation of the endothelial-like tubular structure. Furthermore, we identify distinct Ca^2+^ activity profiles during the different morphogenetic events of notochord development. By employing cell type-specific optogenetics^83^, we show that sequestration of Ca^2+^ from notochord cells prior to and during convergent extension results in defective cell intercalation. Using pharmacological perturbations in combination with Ca^2+^ imaging and quantitative analysis of cytoskeletal and cell junction marker dynamics, we show that gap junctions, stretch-activated ion channels (SACs)^84^, inositol triphosphate receptor (IP3R) signalling, Store Operated Calcium Entry (SOCE) and Sarco/endoplasmic reticulum Ca^2+^-ATPase (SERCA) are major modulators of Ca^2+^ dynamics during tubulogenesis. Finally, we found that Ca^2+^ plays a role in actomyosin regulation *in vivo.* Our findings provide an entry point into dissecting the mechanistic underpinnings of how Ca^2+^ signalling regulates the development of a tubular organ.

## Results

### Quantifying calcium dynamics in the developing notochord

To investigate calcium (Ca^2+^) signalling dynamics in the developing notochord, we used the genetically encoded calcium indicator (GECI) GCaMP6s^85^. GCaMP6s were expressed using the regulatory elements of the notochord specific genes Brachyury^63^ and carbonic anhydrase 3^86^.

Strikingly, we detected spontaneous Ca^2+^ activity in the notochord throughout the developmental stages (stg17-stg26) and morphogenetic events that we imaged (Figure1A-B). Qualitative observation of the recorded Ca^2+^ activity suggests developmental stage-dependent differences (Figure1B). To explore this possibility in a quantitative manner, we first compared Ca^2+^ peak features across the different developmental stages (Figure1C, Sup. Fig.1A, B). We quantified parameters such as peak amplitude, peak base duration (distance between the left and right base), rising and falling slopes (slope of the line drawn from the left and the right base to the peak respectively) and area under the peak. These peak features changed in a dynamic manner across stages, showing stage-specific signatures. Peak amplitude, area under the peak and peak base duration values increased during notochord convergent extension up to the transition from intercalation to cell elongation (stg 17 → stg 21). These features subsequently decreased during cell elongation (stg 22 → stg24), with a further drop occurring during tubulogenesis. Focusing on the changes in falling slope of Ca^2+^ transients across notochord development revealed two distinct phases, one covering cell intercalation and the onset of cell elongation (stg 17 → stg 22) and another corresponding to the late phases of cell elongation and lumen formation (stg 23 → stg 26). On the other hand, rising slope data show that Ca^2+^ peaks have a slower rise during cell intercalation while progressively becoming faster during cell elongation, and significantly faster during the tubulogenesis phase.

Our peak features analysis indicates that notochord cells exhibit Ca^2+^ transients that show differences depending on which morphogenetic stage they are in (cell intercalation, cell elongation or lumen formation). However, as it is often the case with developmental systems that exhibit spatiotemporally complex and irregular Ca^2+^ dynamics, peak features alone have relatively limited resolving power. Dynamic Ca^2+^ signalling at the cellular level is a result of interlinked molecular modules that adjust their behaviour in response to internal and environmental cues. Differential activation of individual modules could trigger a sequence of specific Ca^2+^ signal shapes, generating a unique Ca^2+^ profile. To overcome the limits of conventional methods (primarily in their resolving power), we employed a contemporary machine learning-based analysis method based on time series clustering, to reveal biologically distinct Ca^2+^ dynamics across notochord development. Among numerous existing approaches for time-series clustering^87^, we selected k-Shape clustering^88–90^, which is a recently developed clustering technique that outperforms most other clustering techniques in terms of shape-based time series clustering^88^. K-Shape clustering would allow us to compare on an entire shape-by-by basis, rather than individual peak-features. k-Shape based analysis can thus be particularly useful when dealing with complex multicellular biological systems, such as the developing notochord, which we find to display rich and dynamically changing Ca^2+^ activity. When additionally combined with methods such as Markov Chains, k-Shape based analysis will be able to identify Ca^2+^ activity state changes that are likely also linked to developmentally relevant cell state changes. We thus performed k-Shape clustering analysis of our entire notochord Ca^2+^ imaging dataset, classifying it into 11 clusters (Figure1D). For each cluster, the mean trace was assigned an archetype-alphabetical label according to its half peak width (A → K; arranged from narrower to wider peaks) (Figure1E). We then quantified the contribution of each k-Shape cluster to the Ca^2+^ activity in each developmental stage (Figure 1F; Sup. Fig.1J). During cell intercalation and elongation stages, we found an enrichment of wider peaks. In particular, during stages 17 and 18, all clusters were represented with a small enrichment of cluster I (stg 17: 21.1%, stg 18: 17.0%). Stages 19 to 23 show a progressive increase in the contribution of the clusters with the widest half peak widths; namely clusters K and J, are overrepresented (Figure1F).

Interestingly, stage 21 shows a very large representation of cluster K (50.0%) whereas other clusters such as B, C, E, H & G are absent during that stage, though we note that embryos during this stage show reduced Ca^2+^ activity and thus we have a smaller number of traces contributing to k-Shape analysis for stage 21. During later developmental stages associated with tubulogenesis, stage 24 shows an intermediate signature with decreasing percentages of I (14.1%), J (21.2%) and K (42.4%) archetypes together with a concomitant increase of cluster A (29.5%). Stages 25 and 26, which are associated with the fusion of the lumen pockets and bidirectional crawling movements of notochord cells, show an enrichment in the sharper peaks with shorter half-peak widths especially in the case of clusters D and E. In summary, k-Shape cluster distribution changes across developmental stages allowed us to distinguish between biologically distinct phenomena, such as cell intercalation, cell elongation and lumen formation.

Having established that each of these archetypical peaks is present to a different extent in the developmental stages and morphogenetic processes that notochord cells undergo, we asked whether there are discernible structures in the Ca^2+^ activities across development. Using the assigned alphabets of archetypical peaks obtained through k-Shape clustering (Figure1D,E), we reduced our Ca^2+^ imaging traces to sequences of discrete letters that we subsequently modelled using a statistical Markov Chain model^91^. To visualize the model, we generated a Markov Chain graph (Figure1G; Sup. Fig.1C-I)), a State Transition Matrix of Markov Chain (Figure1H; Sup. Fig1 C-I) and Markov Chain model-based Ca^2+^ traces for each developmental stage (Figure1I; Sup. Fig 1 C-I). Interestingly, all three visualizations show that stages 17, 18, 25 and 26, which are characterized by high cellular motility, exhibit qualitatively more complex transition patterns between the archetypical peaks, while the remaining stages show simpler transition patterns. We defined two measures called *Forward Bias* and *Reverse Bias* based on the State Transition Matrix for each developmental stage. The *Forward Bias* is defined as the sum of the upper triangle of the State Transition Matrix, while the *Reverse Bias* is defined as the sum of the lower triangle of the State Transition Matrix. Values along the diagonal were excluded for both measures. A high *Forward Bias* value represents the states in these traces that tend to transition from lower states to a higher states, with an ascending direction in the archetype-alphabet (e.g. A → D) where shorter peaks are more likely to be followed by broader peaks. In contrast, a high *Reverse Bias* value indicates that the states within these traces tend to transition from higher states to lower states (i.e. in a descending direction in the alphabet, e.g. D → A). The majority of the State Transition Matrices show a higher *Forward Bias,* with stages 19 and 22 showing the largest difference between *Forward* and *Reverse Biases,* suggesting a “locking” of Ca^2+^ signalling patterns which have a high tendency to produce sharper peaks followed by broader peaks (Sup. Table 11). Only 2 out of 10 developmental stages have a higher *Reverse Bias,* namely stage 24 and stage 25 (Sup. Table 11). The model-based Ca^2+^ traces (Figure1G-I) were qualitatively similar to the real notochord cellular Ca^2+^ activity we recorded (Figure1B). This suggests that spontaneous Ca^2+^ activity observed in the notochord can be captured using comparably simple stochastic models. Both the modelled and real observations are in agreement with a progressive increase in the representation of wider peaks from stage 19 until stage 23 as discussed earlier, and the gradual enrichment in sharper peaks at later developmental stages (stages24-26).

To summarize this part, our peak feature analysis suggests that the morphogenetic processes taking place during notochord development are characterised by Ca^2+^ transients whose visually distinguishable features (e.g. amplitude and area under the curve) dynamically change across development. Using k-Shape clustering and Markov Chains, we mapped Ca^2+^ signalling complexity across notochord development with our data indicating that Ca^2+^ signalling shows plasticity in the organization of the Ca^2+^ signals. These changes across development may orchestrate cell behaviour changes.

### Optogenetic sequestration of calcium reveals the requirement of calcium signalling in notochord convergent extension

Having established the presence of Ca^2+^ activity in the notochord across developmental stages, we next addressed the role of Ca^2+^ signalling in development. To this aim, we manipulated Ca^2+^ levels on demand and in a cell-specific manner in the notochord cell cytoplasm using an optogenetic actuator, PACR^83^. PACR is a tool that combines a photosensitive protein domain, LOV2, and a Ca^2+^ binding protein to enable Ca^2+^ release in a light-inducible manner. In the absence of blue light, Ca^2+^ is bound to PACR and thus causes Ca^2+^ to be sequestered from the cytoplasm. Upon activation with blue light, Ca^2+^ is released by PACR into the cytoplasm. Embryos expressing PACR in the notochord under the control of the Brachyury promoter^63^ were split into two groups. One group was kept in the dark while the other was continuously exposed to blue light during development (Figure2A). Tailbud embryos were then fixed at stage 23/24. Embryos kept in the dark were on average shorter compared to embryos exposed to blue light and, remarkably, had irregular arrangement of their notochord cells along the A-P axis. These suggest that embryos kept in the dark were unable to complete notochord convergent extension. To quantify the morphogenetic defects, we labelled the Brachyury>msEGFP-PACR embryos with phalloidin and performed 3D confocal imaging. We then mapped the outline of the embryo and the centroids of all notochord cells in the tailbud embryos (Figure 2A, B), and found that embryos kept in the dark have a shorter tail compared to those illuminated with blue light. We next quantified the extent of intercalation defects by measuring the distance of each notochord centroid to the embryo midline and the intercellular distance between adjacent notochord cells. We demonstrate that PACR embryos exposed to blue light show little variation in the distance to skeleton, while those kept in the dark have an increased distance between notochord cells and skeleton (Figure2D). These data indicate that embryos where cytoplasmic Ca^2+^ is sequestered by PACR have reduced ability to perform cell intercalation. In summary, our optogenetics experiments demonstrate that sequestering Ca^2+^ results in reduced fidelity of the convergent extension process.

**Figure 2:**
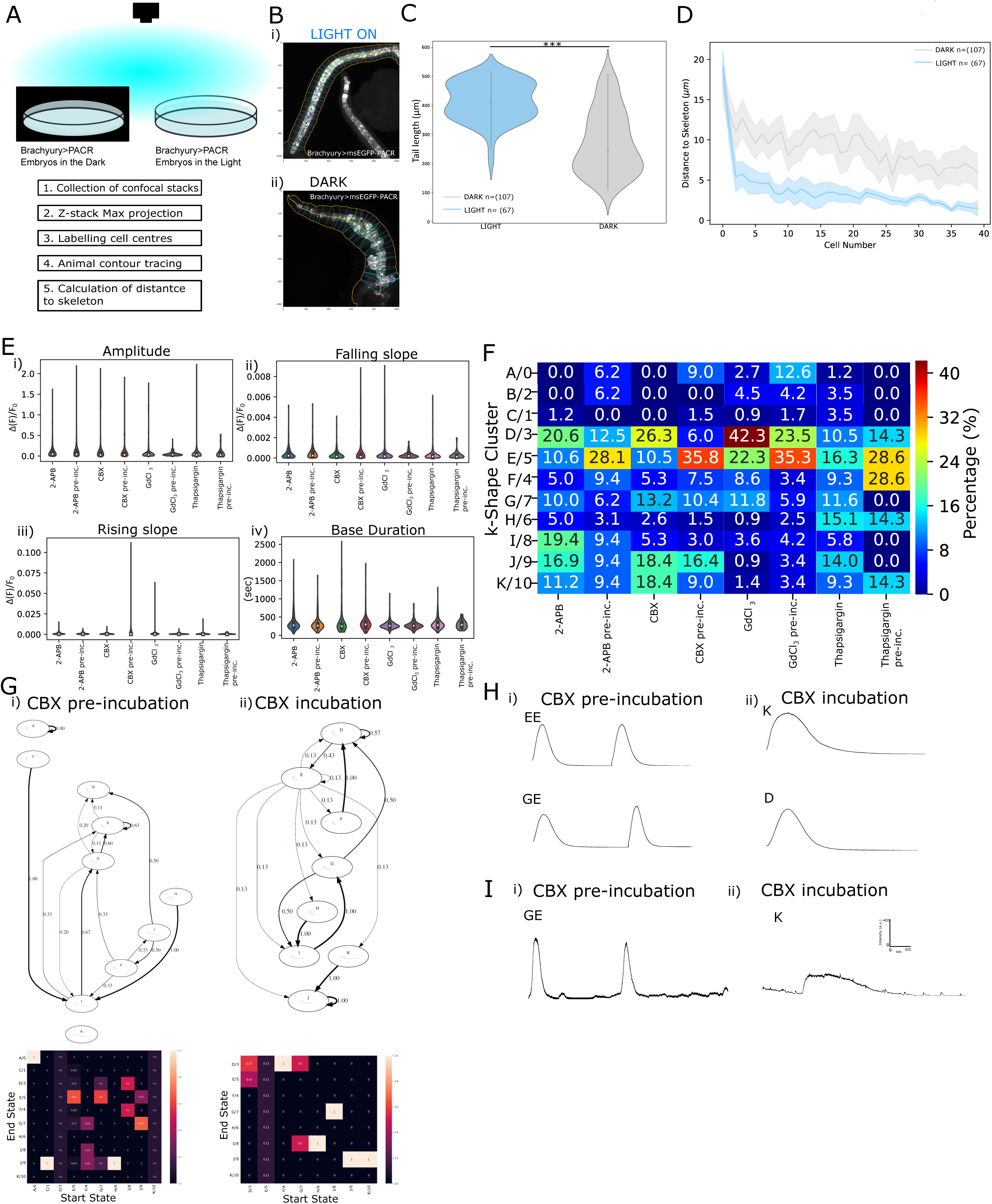
Ca^2+^ signalling is required for cell intercalation, while Ca^2+^ activity depends on SACs, IP3R, SOCE, ER-stored calcium and gap junction coupling. A) i)Schematic of optogenetic assay. Embryos electroporated with the optogenetic actuator Brachyury>PACR were divided into two groups. One group was shielded from blue light while the other group was exposed to blue light. A stepwise overview of the analysis pipeline. B) Maximal projections of two embryos from the i) illuminated with blue light, ii) kept in the dark showing body contour (orange), notochord skeleton (light green), cell centroids are marked with a coloured dot. The blue dashed lines show the distance of the centroid from the embryo contour. C) Quantification of notochord skeleton length in μm, Mann-WhitneyU test *** p= 6.9208820458255084e-15. The number of embryos analyzed for DARK=107 and for LIGHT=67. D) Distance of the notochord cell centroid from the notochord skeleton as a function of the notochord cell position. Cell 0 is the most anterior notochord cell and cell 39 is referring to the most posterior notochord cell. Statistical significance was calculated using Mann-Whitney test with a multiple comparisons Bonferonni correction. Positions 0 and 1 were not significant (p>0.05) while all other positions were (p<0.05) (see Sup. Table22 for p-values per cell position). E) Peak features of Ca^2+^ transients from control and drug treated notochord cells during tubulogenesis stages. 4-18 embryos were used per treatment (see Sup. Table2). F) k-Shape cluster composition (unit is %) of control and drug treated notochord Ca^2+2+^ transients. G) Markov-chain generated transition graph and transition matrix from i) pre-CBX incubation data, ii) Cbx incubation data. H) Example traces generated from the Markov Chain model for pre-CBX (EE, GE) incubation and CBX incubation (K, G). I) Raw Ca^2+^ activity traces from notochord cells before incubation with CBX and during incubation with CBX. The traces are annotated with the corresponding k-Shape cluster letter combination.

### Calcium peak features depend on SACs, while calcium profile organization and composition are additionally dependent on IP3R, SOCE and ER-stored calcium mechanisms

To gain insights into the molecular pathways controlling Ca^2+^ signalling in the developing notochord, we pharmacologically interfered with inositol triphosphate receptor (IP3R) signalling, Store Operated Calcium Entry (SOCE), ER-calcium stores and stretch-activated ion channels (SACs). We blocked IP3R and SOCE signalling using 2-aminoethoxy-diphenyl-borate (2-APB)^37, 92–94^. Embryos from the tubulogenesis stages were incubated for 5 minutes in control buffer and subsequently for 15 minutes with drug. We quantified peak features of Ca^2+^ transients before and during drug treatment (Figure2E; Sup. Fig2 A,B). Surprisingly, 2-APB had modest effects on peak features that were not statistically significant when comparing to controls. However, when we calculated the percentage distribution of k-Shape clusters, we found that 2-APB incubation increased the representation of clusters I (x2.12-fold), J (x1.8-fold), K (x1.2-fold) while decreasing the contribution of clusters A and B. Furthermore, Markov Chain modelling of 2-APB data demonstrated an increase in *Forward bias* (Sup. Figure 2C,E; Sup. Table12). We then tested the contribution of ER-stored Ca^2+^ in modulating Ca^2+^ peak features and activity profile in notochord cells. To this aim, we pharmacologically inhibited the SERCA pumps using Thapsigargin^95–99^. Notochord Ca^2+^ transients were shown not to be sensitive to Thapsigargin treatment when peak features were compared between control and drug-treated conditions, even though as expected, the amplitudes of Ca^2+^ transients were higher in the drug-treated condition compared to controls (Figure2E; Sup. Fig2 A,B). Despite the limited effects on Ca^2+^ peak features Thapsigargin treatment resulted in substantial changes in k-Shape cluster distribution relative to controls. In particular, clusters D (ca 25%), F (ca67%) with a parallel large increase in the contribution of clusters G, I and J. Finally, Markov Chains modelling of the Thapsigargin-treated data revealed an increase in *Forward bias* compared to controls (Sup. Figure 2C,E; Sup. Table12). These suggest that the basic peak features of Ca^2+^ transients are not under the control of IP3R, SOCE-dependent and ER-stored Ca^2+^ mechanisms, yet these pathways do have an influence on the organization and composition of the Ca^2+^ profile of notochord cells during tubulogenesis, the latter finding made only possible with our new analytical methodologies.

During embryonic development, cells and tissues generate mechanical forces. The effects of these forces on morphogenetic processes have been linked to Ca^2+^ signalling^100, 101^. In addition, the developing notochord of *Ciona* is known to express ion channels that may transduce mechanical forces into Ca^2+^ signals^62^. Thus, to test whether SACs are involved in the generation of Ca^2+^ activities during tubulogenesis, we applied a pharmacological inhibitor, GdCh^102, 103^. This revealed that GdCh strongly affects all features besides duration base. Calculating the percentage distribution of k-Shape clusters demonstrated that GdCh treatment results in almost two folds increase of clusters D, F and G compared to controls (Figure2F; Sup.Fig2F). Modelling the control and drug-treated data with Markov Chains further revealed that GdCh treatment results in an overall *Reverse bias* (Sup. Fig.2D; Sup. Table12). Our findings suggest that SACs are important contributors to all aspects of Ca^2+^ activity recorded by us during notochord tubulogenesis.

### Dissecting the role of intercellular calcium signaling through gap junctions during tubulogenesis

We next addressed cell-cell Ca^2+^ signalling. A common way of transmitting intercellular Ca^2+^ waves in non-excitable cells like notochord cells is through the exchange of Ca^2+^ or IP3 via gap junctions^104^. During tubulogenesis, a series of coordinated events between neighbouring cells are required to take place. It is conceivable that intercellular communication plays a role in maintaining and coordinating Ca^2+^ dynamics during notochord morphogenesis. To test this hypothesis, we applied a gap junction inhibitor, CBX^105–107^. Incubation with CBX decreased Ca^2^+ dynamics compared to controls, as shown by significantly reducing most major peak features including amplitude, base duration, falling and rising slopes. When k-Shape clustering analysis was applied, we found that CBX treatment results in a decrease in clusters A and E, a large increase in cluster D, and a smaller increase in cluster K (Figure2F). The directionality of the State Transition Matrix for the CBX data showed a switch to an overall *Reverse bias* compared to control data (Figure2F), indicating a switch in the composition and organization of Ca^2+^ activity in individual notochord cells in the absence of functional cell-cell coupling. Combined, our data demonstrate that Ca^2+^ signalling in notochord cells requires gap junction mediated cell-cell coupling.

### Calcium signalling regulates remodelling of the cytoskeleton and the localization domains of cell adhesion complexes

Having demonstrated by k-Shape clustering and Markov Chain modelling methods that the developing notochord of Ciona embryos exhibits complex Ca^2+^ activity, we addressed its relevance in physiology. Deciphering the molecular and cellular effectors of Ca^2+^ signalling *in vivo* remains a challenge. Amongst them, cytoskeletal^108, 109^ and cell-cell adhesion^110–112^ effectors have been shown to mediate subcellular functions of Ca^2+^ signalling that have implications in both normal and pathological contexts. We thus visualized the cytoskeleton using fluorescently tagged markers -for the actomyosin network, Actin^77^, Utrophin and Myosin Light Chain (MLC)^76, 77^; and the microtubule network, Ensconsin^77^. We performed two types of pharmacological perturbations, one long-term (late gastrula until stage 24) and one acute (stage 24 only), using in both cases drugs 2-APB, GdCh and CBX. To quantify the morphological changes of the cytoskeleton under the different pharmacological perturbations, we employed the box-counting method that enables analysis of complex patterns. As a result of this analysis, we obtained a quantitative metric termed fractal dimension, a statistic that has been previously employed to analyze both two-dimensional and 3D cytoskeletal structures in tissue culture cells^113–117^. We applied fractal dimension to quantitatively describe the complexity of our 3D *in vivo* data. Here, a decrease in fractal dimension values indicates a decrease in network complexity and vice versa; even though decreases and increases in complexity do not directly translate to loss-of-function and gain-of-function respectively.

Imaging of Actin revealed that long-term treatment with CBX and GdCh resulted in significantly lower fractal dimension values relative to control, while acute treatment with 2-APB and CBX gave rise to a significant decrease in fractal dimension values relative to control (Figure3A). Long-term treatment with CBX and GdCh respectively decreased and increased the fractal dimensions of Utrophin, while all acute treatments resulted in a strong decrease in fractal dimensions of Utrophin (Figure3B). The final actomyosin marker that we tested, MLC, showed no change in fractal dimensions under all long-term drug treatments, while acute treatments with both 2-APB and CBX resulted in increases in fractal dimensions of MLC (Figure3C). In contrast, the microtubule marker Ensconsin showed only a significant increase in fractal dimension upon long-term incubation with GdCl_3_, and no statistically significant change under acute drug treatments even though we observed a trend for decreasing fractal dimension for all drug treatments (Figure3D). In summary, these data suggest that Ca^2+^ signalling is required for actomyosin network organization, but may be dispensable for regulation of the microtubule network organization.

**Figure 3:**
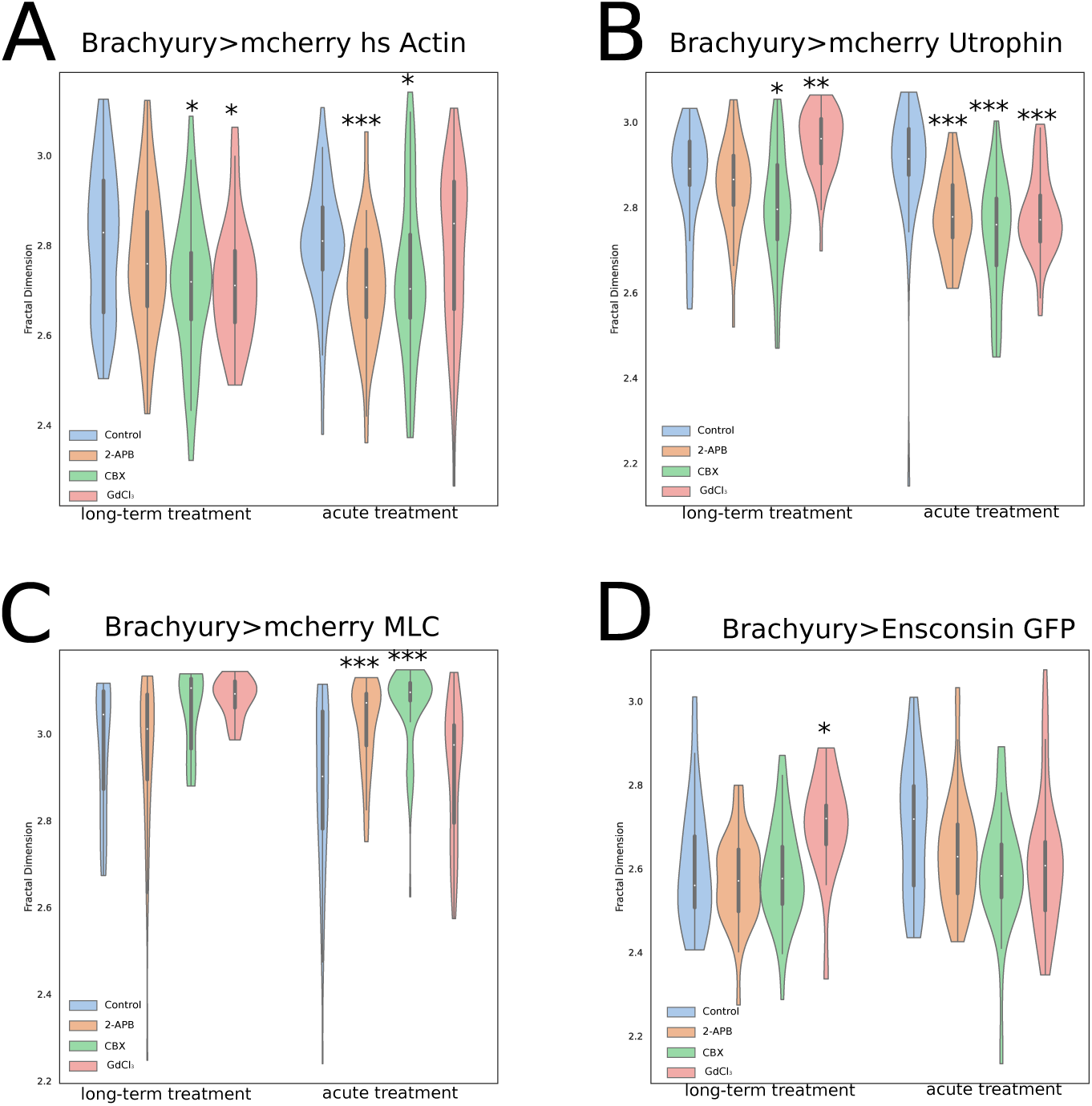
Ca^2+^ signalling affects actomyosin organization. Quantification of cytoskeletal markers in control and long-term or acutely treated embryos with CBX, GdCl_3_ and 2-APB. (A-D) Violin plots of Fractal dimension values using the box counting method, Minkowski-Bouligand dimension for transgenic embryos harbouring assayed transgene. Statistical significance was calculated using the Mann-Whitney U-test with Holm correction (*0.05<p, **0.005<p, ***0.0005<p) (p-values can be found in Sup. Tables4-7) (A) Brachyury>mCherry-hsActin A) Brachyury>mCherry-Utrophin, C) Brachyury>mCherry-MLC, D) Brachyury>Ensconsin (see Sup. Table 3 for numbers of animals used).

Having established that Ca^2+^ signalling regulates remodelling of the actin network during notochord tubulogenesis, we next probed for interdependence between the two processes. Specifically, we asked whether disruption of actin polymerization alters Ca^2+^ signalling dynamics. To this aim, we applied a recently developed tool, DeAct-SpvB^118^ that expresses a mono(ADP-ribosyl)transferase domain, which renders actin monomers unable to polymerize. We drove expression of DeActSpvB under the cah3 promoter and recorded Ca^2+^ activity in the notochord during tubulogenesis stages (stages 24-26) using Brachyury> GCaMP6s. DeAct-SpvB embryos’ Ca^2+^ activity showed a higher frequency of Ca^2+^ transients. (Figure4A-C). Analysis of Ca^2+^ transient peak features revealed that disrupting actin dynamics significantly reduced all Ca^2+^ peak features (Figure4D). We then analyzed the k-Shape cluster distribution between control and DeAct-SpvB embryos, and found that the contribution of most clusters decreased with a concomitant large increase (∼2x) for clusters H, G and I. State Transition Matrices for control and DeAct-SpvB data show an overall increase in Forward Bias (Figure5F-G; Sup. Table13). Our data demonstrate that a functional actin cytoskeleton is required to maintain wild-type Ca^2+^ dynamics during tubulogenesis and that blocking intracellular and intercellular Ca^2+^ signalling alters the complexity of the cytoskeletal network. Taken together our findings indicate existence of a feedback mechanism between Ca^2+^ activity dynamics and actin cytoskeleton organization in the developing notochord.

**Figure 4:**
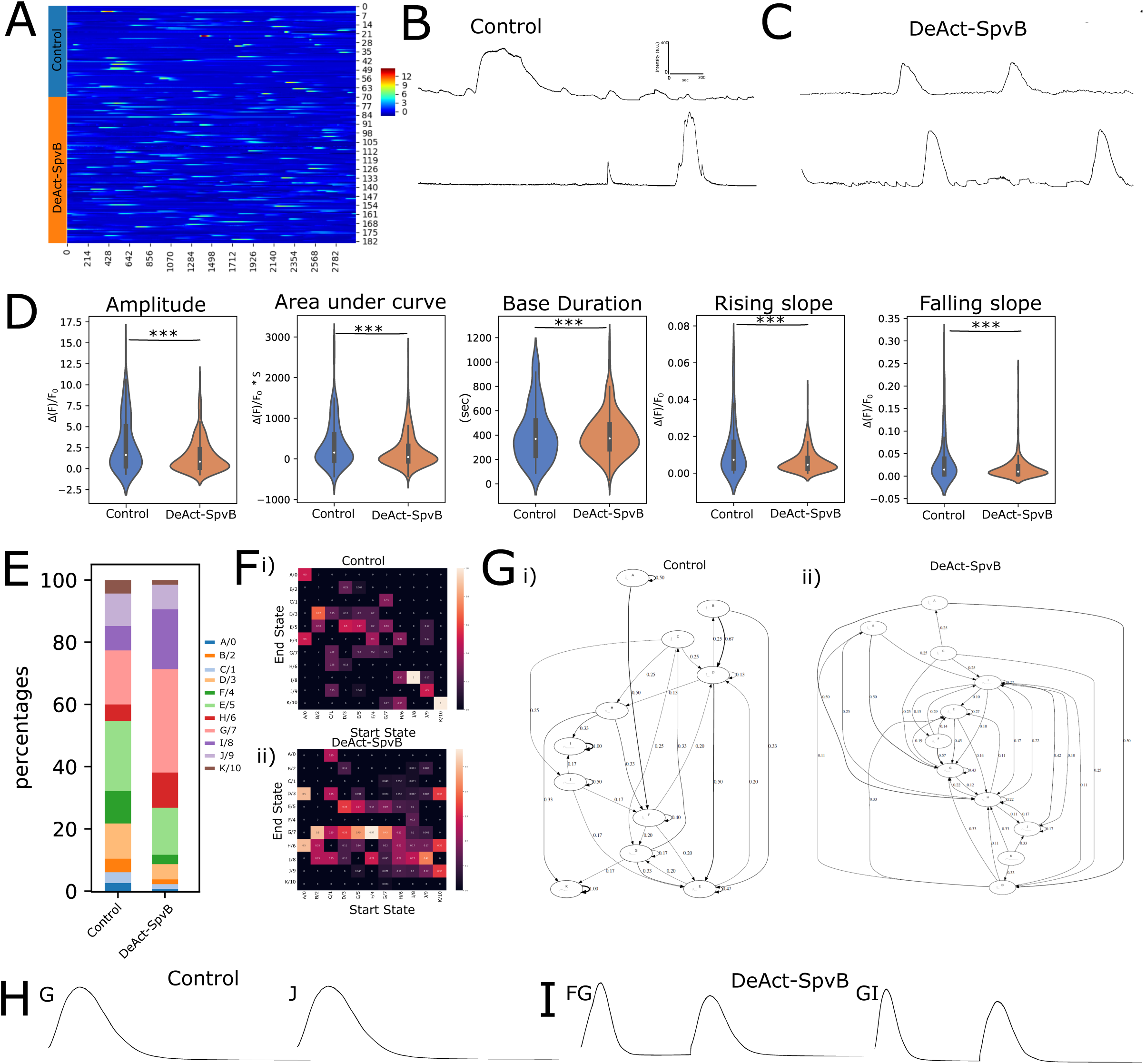
Actin depolymerisation alters Ca^2+^ dynamics during tubulogenesis. (A) Amplitude based heat map for control and cah3>DeAct-SpvB Ca^2+^ data. (B-C) Example traces from Control and DeAct-SpvB expressing notochord cells. (D) Quantification of Ca^2+^ peak features of control (n=24) and cah3>DeAct-SpvB (n=32) expressing embryos (Sup. Table 8). E) k-Shape cluster distribution in control and cah3>DeAct-SpvB embryos. F-G) Markov-Chain model generated transition matrices and transition graphs for i) Control and ii) cah3>DeAct-SpvB. Example traces generated from the Markov Chain model for control (H) and cah-3>DeAct-SpvB (I).

Finally, to determine whether the above pharmacological perturbations of different aspects of Ca^2+^ signalling had any effects on the integrity of cell junctions, we labelled the adherens junctions with E-cadherin^74^ (Figure5A). We show that E-cadherin in control embryos occupies the lateral domains (the anterior and posterior surfaces) of notochord cells (Figure5C), as previously described^74^. Surprisingly, embryos treated with GdCl_3_, CBX and 2-APB at late gastrula stage show altered E-cadherin distribution, with strong levels of expression not only located on the anterior and posterior sides of notochord cells but also on their basal surface (Figure5C). To quantify the effects of the different treatments, we compared the peak intensity ratios generated using the fluorescence intensity profiles of the lateral and basal surfaces of notochord cells (Figure 5B,D). All treatments led to a decrease in lateral versus basal peak ratio relative to the control. The strongest effects were observed in treatments with 2-APB and GdCh, while CBX treatment gave rise to a moderate drop (Figure 5D). Our data suggest that intracellular Ca^2+^ signalling via the IP3R and SOCE pathways, and to a lesser extent intercellular communication via gap junctions, are required for the correct localization of cell junctions along the lateral domain during early phases of tubulogenesis.

**Figure 5:**
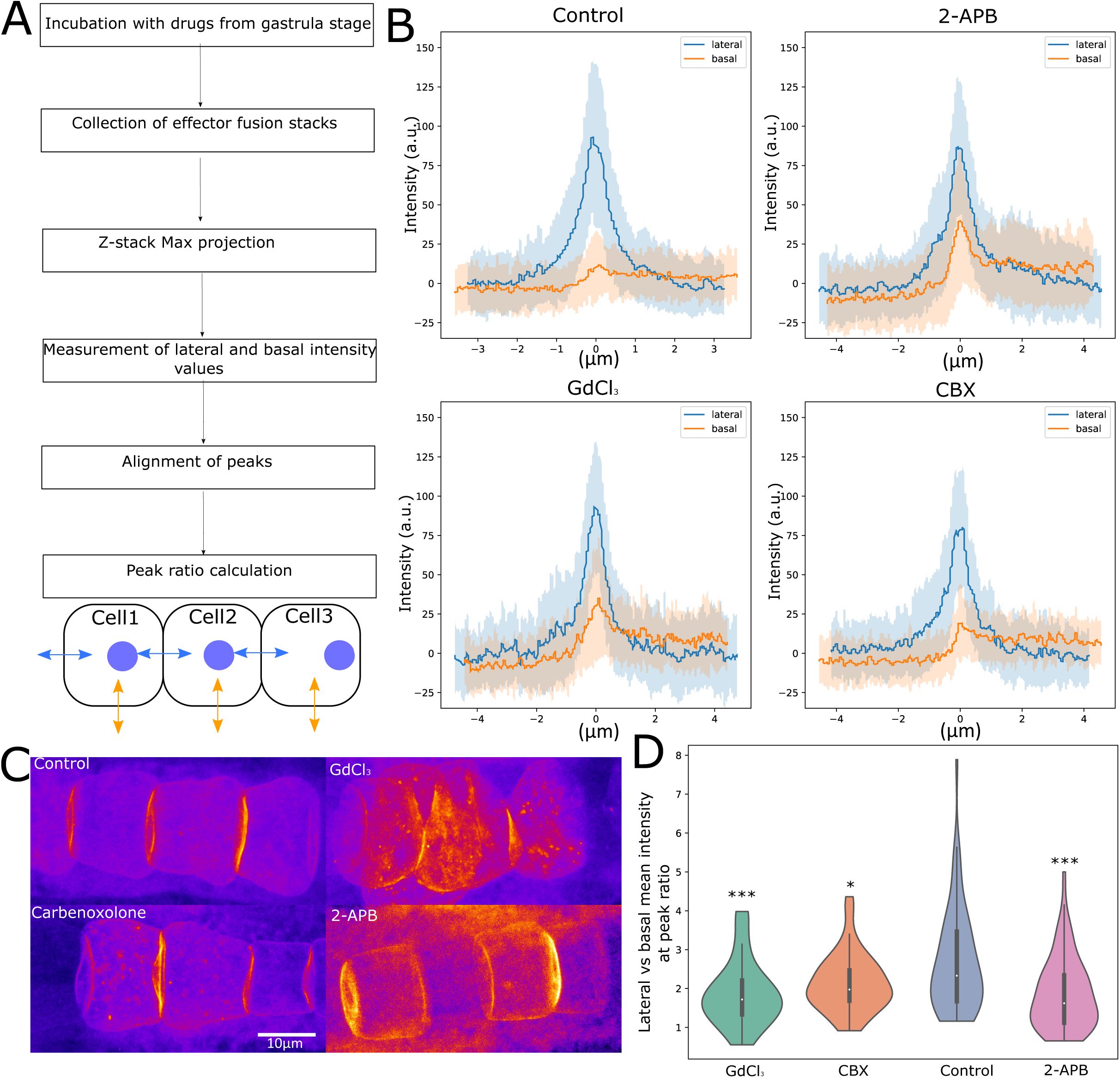
Ca^2+^ signalling limits E-cadherin expression to the apical domain of notochord cells. A) Pipeline for determining the Dm-E-cadherin lateral vs basal mean intensity peak ratio. B) Mean traces of fluorescent intensity profiles for control, CBX, GdCl_3_ and 2-APB. Lateral intensity profile shown in blue, basal intensity profile shown in orange. Blue and orange shades indicate S.E.M. (see Sup. Table 9 for number of measurements per condition). C) Representative pseudocoloured images showing maximal projections of confocal stacks from notochord cells expressing Brachyury>Dm-E-Cadherin-mCherry. D) Violin plots quantifying the mean intensity at peak ratio between the lateral and basal expression domains of Dm-E-Cadherin-mCherry. Statistics were calculated using Mann-Whitney test * p<0.05, *** p<0.0005. (detailed p values in Sup. Table 10)

## Discussion

Here we recorded and quantified Ca^2+^ dynamics throughout the embryonic development of a tubular organ present in all chordates, the notochord. We demonstrate the presence of Ca^2+^ activity during multiple critical morphogenetic events that are commonly employed by developing tissues and organs across the animal kingdom, including cell intercalation, cell elongation and tube formation. By performing *in vivo* live imaging, we found that notochord cells alter the properties (peak features) of their Ca^2+^ transients in a dynamic manner according to the developmental stage. Furthermore, k-Shape clustering has enabled us to generate a ‘Ca^2+^ curve shape-based alphabet’ system that was applied to our time-series data, revealing that such archetypical ‘peaks-alphabet’ is employed in a specific manner according to real time cellular behaviours. Markov-Chain based modelling uncovered that the sequence-structure of Ca^2+^ signalling peaks changes with developmental stage. Moving on to targeted perturbations, we employed an optogenetic tool, PACR, in a spatiotemporally defined manner to demonstrate that Ca^2+^ signalling is essential for the successful completion of cell intercalation. In addition, pharmacological perturbations combined with Ca^2+^ imaging revealed that IP3, SOCE and SERCA-based Ca^2+^ signalling components are involved in notochord tubulogenesis. By perturbing gap junction function, we uncovered a higher variability in intracellular Ca^2+^ oscillations in embryos that have disrupted intercellular connections. This is consistent with previous reports that gap junction-mediated intercellular communication exists in multiple non-excitable cell types^119–121^ and tissues^107^.

At the tissue level, it has been suggested to confer uniform response to a localized cue such as metabolic activity^120^, vindicating the use of multicellular systems to study dynamic calcium signalling. Our study is the first to perform a quantitative characterization of the dynamic relationship between Ca^2+^ signalling and its cellular effectors, including cytoskeletal and cell junction components, in the context of the developing notochord. We combined quantitative imaging of well-characterized cytoskeletal and adherens junction markers with pharmacological inhibitors of different aspects of Ca^2+^ signalling regulation. Quantitative analysis allowed us to assess differences in actomyosin and microtubule network complexity between control and drug-treated notochord cells. Pharmacological perturbations, particularly the acute treatments, significantly altered the complexity of the actomyosin network, while effects on the microtubule network were extremely modest, suggesting that Ca^2+^ signalling regulates actin cytoskeleton but not microtubules in the *Ciona* developing notochord. Interestingly, Ca^2+^ signalling has recently been linked to actomyosin reorganization and activation during development^107^, in response to wounding induced damage^122, 123^, wound healing^124, 125^, tissue regeneration^125, 126^, and cell extrusion^127, 128^. Furthermore, Ca^2+^ signalling has been shown to be directly coupled to actin dynamics during tip growth in the protonema of *Physcomitrella patens*^129^.

This work has also uncovered a feedback mechanism between Ca^2+^ signalling and the actin cytoskeleton: we show that not only does Ca^2+^ signalling regulate organization of the actin cytoskeleton, but blocking actin polymerization also heavily affects Ca^2+^ peak features, and results in substantially altered k-Shapes derived clusters distribution. It has previously been suggested that the actin cytoskeleton can influence Ca^2+^ activity in plant cells^130^, starfish oocytes^131, 132^, and hippocampal^133^ neurons^133^.

In our study, pharmacological perturbation of different aspects of Ca^2+^ signalling leads to ectopic expression of E-cadherin on the lateral side of the notochord cells. Changes in cell-cell contacts likely result in changes in cell shape, as well as in the mechanical and signalling properties of the cells.

Tube formation is fundamental to the development of functional organisms. Nevertheless, there have been relatively few studies that have addressed the role of Ca^2+^ signalling in this process. Two studies especially relevant to this, which have performed *in vivo* and *ex vivo* Ca^2+^ imaging respectively during tube formation and differentiation, have shown that Ca^2+^ signalling is necessary for neural tube closure in ascidians and Xenopus^134^ and pronephric tubule differentiation in Xenopus^43^. A more recent study characterizing Ca^2+^ dynamics in *Ciona robusta* and *Ciona savignyi* during early embryogenesis has reported Ca^2+^ activity in the neural tube, and additionally identified Ca^2+^ activity in the notochord during cell intercalation^33^. Here we have discovered that Ca^2+^ activity in the *Ciona intestinalis* notochord is not limited to cell intercalation as previously reported, but instead is present throughout notochord development, including the notochord elongation and tubulogenesis phases. What might be the role of Ca^2+^ activity in these morphogenetic processes during notochord development in *C. intestinalis?* Our analysis revealed that the different steps in the morphogenesis of the notochord coincide with changes in the profile of Ca^2+^ activity. Our data raise the possibility that the shape and temporal dynamics of Ca^2+^ fluctuations are robustly encoded into specific cell behaviours. Furthermore, *C. intestinalis* notochord has been reported to express several Ca^2+^ signalling transducers, including CaMK, MLCK, Calumenin and the transcription factor NFAT5^59, 61, 62, 135, 136^. Therefore Ca^2+^ activity in the notochord likely regulates cell behaviour through molecules that regulate various types of outputs ranging from gene expression to cell shape and cell motility. Methodologically, our work presents broadly applicable advances in two directions. Firstly, we employed a recently introduced machine learning based analysis method, k-Shape clustering^88^, with which we have been able to quantitatively describe the heterogeneity, hierarchy and structure of notochord Ca^2+^ dynamics throughout development. Indeed, several non-excitable cell types exhibit heterogeneous Ca^2+^ activity^20, 33, 41, 137^–^139^. Early studies employing quantitative Ca^2+^ imaging had generated hierarchies based on the signal initiation site and the kinetics of the observed activity ^137, 140, 141^. Here we used a semi-supervised method to capture in 11 clusters the vast majority of the heterogeneity of our entire dataset. Using k-Shape clustering analysis, we identified developmental stage-specific differences in the heterogeneous usage of these clusters. In addition, using Markov Chains^91^, we were able to exploit the k-Shape derived clusters to investigate the underlying syntax (temporal organization) of the Ca^2+^ activity we recorded.

A further methodological advancement is the use of two experimental tools, which to our knowledge is the first time to be applied in the context of a developing tubular organ and in non-excitable cells. The first one, PACR, belongs to a family of optically-controlled Ca^2+^ actuators, which enabled us to demonstrate that Ca^2+^ signalling is required for successful completion of convergent extension. The second tool is DeActs-SpvB, which allowed us to, in a cell specific manner, address the potential feedback between actin cytoskeleton organization and Ca^2+^ activity dynamics. We believe that future studies will greatly benefit from tools that can alter signalling and/or cytoskeletal dynamics in a cell-specific and temporally controlled manner with even higher precision and reversibility. We show that longer-term pharmacological treatment gives milder phenotypes in cytoskeletal organization. Interestingly, this raises the possibility that the notochord and the associated Ca^2+^ signalling might be subject to a form of developmental adaptation. This further emphasizes the requirement for acute perturbations combined with real-time recording of behaviour when studying systems that can adapt^142^.

In our study we have laid the foundations for using the notochord of *Ciona* as a model for a systems level dissection of the contribution of Ca^2+^ signalling to the morphogenetic processes that shape tubular organs. The recent establishment of cell-type specific CRISPR/Cas9^143^ in *Ciona* raises the possibility of dissecting, in an organ specific manner, the molecular players that encode and decode Ca^2+^ signals and regulate notochord cell behaviour^7, 24, 144^. The presence of the Ca^2+^ responsive transcription factor NFAT5 in the developing notochord of *Ciona* raises the tantalizing possibility that Ca^2+^ dynamically regulates the transcriptional cell state of notochord cells. In this scenario, NFAT5 could be acting as a Ca^2+^ activity frequency decoder as documented in other systems^145–148^. At the same time, further efforts should be invested in understanding the link between Ca^2+^ signalling and actomyosin network changes during notochord formation.

The developing *Ciona* notochord is also an excellent system for applying genetically encoded tools such as optogenetic actuators. These tools will allow us to manipulate Ca^2+^ signalling with high temporal resolution and subcellular accuracy (at the level of organelles). In combination with computational modelling, we can anticipate deciphering general principles of how Ca^2+^ dynamics coordinate morphogenetic processes such as convergent extension and tubulogenesis.

## Methods

### Animals

We collected adult *Ciona intestinalis* from following sites in the Bergen area: Dosjevika (Bildoy Marina AS), Døsjevegen, 5353. The animals were kept in a purpose-made facility, using filtered seawater at 9°C with a pH of 8.2 and a salinity of 3,5%, under constant illumination to stimulate egg production. We obtained eggs and sperm from individual animals to perform in vitro fertilization.

Eggs without chorion were fertilized at the same time and incubated in artificial sea water (ASW, Red Sea Salt) at 14 °C. The ASW pH was 8.4 at 14°C with a salinity around 3,3-3,4%.

### Dechorionation and electroporation procedure

We performed dechorionation according to standard protocols^149^. We collected sperm and eggs from 3-4 adults per experiment. We placed the eggs in dechorionation solution (100mg Sodium Thioglycolate (Sigma T0632), 9mL ASW, 32μ! 10M NaOH, 1mL 0.1% protease E (Sigma P5417)) and agitated gently by pipetting until the follicle cells and chorion were removed. The fertilized, dechorionated eggs were then washed 3 times in ASW before electroporation. We washed the eggs, added the sperm, and left to fertilize for 9 minutes. Electroporation mixes contained a combination of 400 μ! 0.96M mannitol and 100 μ! plasmid DNA were combined in an electroporation cuvette to which 200 mL of fertilized, dechorionated eggs in ASW were added. The cuvette was shocked in a Bio-Rad Gene Pulser XCell using 50 mV, a capacitance of 900-1200μF and a 4mm cuvette gap. We used electroporations that achieved a time constant between 12ms and 40ms.

### Molecular cloning

In order to drive notochord specific expression of various reporters and actuators in the notochord throughout development we generated two promoters: a 1.66kb promoter upstream of the cah-3 (KH.C1.423) gene and a 3.32kb promoter upstream of the Brachyury (KH.S1404.1) gene (see table for primer details). We used BP Clonase II to recombine our PCR product with the P4-P1R. The generated pENTRY clones pMCH1436 (pCah-3) and pMCH1844 (pBrachyury), which were recombined with plasmids pMCH1377 GCaMP6s and pMCH754 unc-54 3’UTR and pDESTII. We used LR Clonase II to perform the recombination reaction. To generate the Entry clone for DeAct-SpvB we subcloned the relevant sequence (1407bp long) from pCMV-DeAct-SpvB. This was a gift from Brad Zuchero (Addgene plasmid # 89446; http://n2t.net/addgene:89446; RRID:Addgene_89446) (please see primer table for details). We used BP Clonase II to recombine the PCR product with PDNOR221. The generated pENTRY clone pMCH2301 was recombined with pCah-3 and unc-54 3’utr using LR Clonase II and pDESTII. To generate the Brachyury>PACR>unc-54 3’UTR construct we amplified the msEGFP-PACR sequence (1752bp long) from msEGFP-PACR_pcDNA3 which was a gift from Takeharu Nagai (Addgene plasmid # 55774; http://n2t.net/addgene:55774; RRID:Addgene_55774). The amplified PCR product was introduced into gateway vector PDNOR221. The pENTRY clone was recombined with the Brachyury promoter (1^st^ position) and the unc-54 3’UTR (3^rd^ position) with pDEST II in order to generate an Expression vector.

Table with primers used in this study:

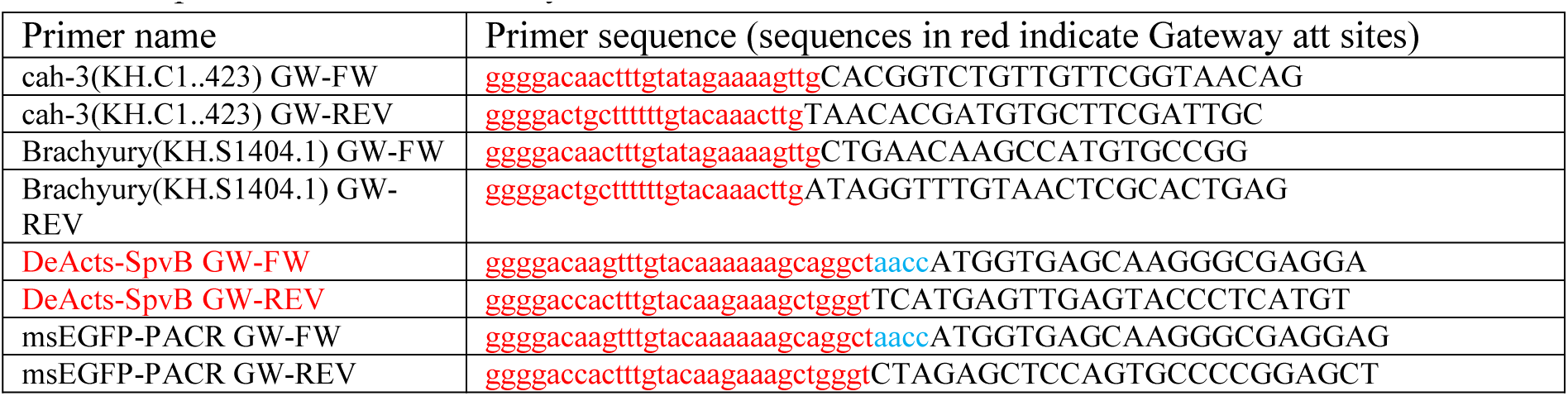

### Calcium imaging acquisition

Electroporated embryos raised at 14°C were imaged at specific developmental stages (17 to 26). Embryos were transferred from their culture plate to a 9cm petri dish in a small drop of ASW. 1.5% low melting point agarose (Fisher BioReagents, BP1360-100) was used to immobilize the embryos on the petri dish. We illuminated embryos using a mercury lamp with a BP470/20, FT493, BP505-530 filterset. Embryos were imaged under a 40x water immersion objective mounted on an Axioskop A1 equipped with a Hamamatsu Orca Flash 4.0V2 CMOS camera. The field of view was 320μm in each direction (2048×2048pixels). Data was acquired using custom made software^150, 151^. 5 minute movies at 10fps were collected. For drug experiments embryos were embedded in 1.5% low melting point agarose in ASW in 9cm petri dishes. Embryos were perfused with control solution for 5 minutes followed by an exchange of the control solution with the drug solution and subsequently we imaged for 15 minutes (3 movies each 5 minutes long). All movies were acquired at 10fps and subsequently analyzed using Mesmerize v0.1^89^ for motion correction, signal extraction, downstream analysis and for producing some of the figures.

### Calcium imaging signal extraction

Raw movies were Motion Corrected using the NoRMCorre^152^ implementation from the CaImAn library^153^ through standard Mesmerize modules. Signal extraction was performed using CNFME^154, 155^ to account for background fluctuations. ROIs with significant spatial overlap or background noise were manually excluded in addition to component filtering performed by CaImAn.

### Downstream calcium imaging analysis

CNMFE-derived traces were normalized within the raw fluorescence values derived from the spatial footprint of the corresponding ROIs. Δ(F)/F_0_ were calculated from raw-normalized traces without using a rolling window. F_0_ was set as the minimum value of the raw-normalized curve. The Butterworth filter was used to smooth the traces and derivatives of the filtered curves were used to find relative maxima and minima which were then placed on the unfiltered curve to avoid filtering-based artifacts^156^. Erroneous peaks and bases were manually removed, and peaks which were clearly undetected were manually added. Peak features were calculated in Mesmerize using z-scored ΔF/Fo values.

Tslearn v0.2.2^90^ was used to perform semi-supervised k-Shape clustering^88^ to class Δ(F)/F_0_ peak curves into 11 clusters. The centroid seeds for k-Shape clustering were obtained in a semi-random manner by first sorting all the curves according to half-peak-width. The sorted curves were then split into 11 equal-sized partitions, and a seed was randomly picked from each partition.

Markov Chains were produced using pomegranate v0.11.2^91^ in a custom-written Jupyter Notebook that can be run as a Voila application (https: //github.com/voila-dashboards/voila): https://github.com/kushalkolar/mesmerize manuscript notebooks/blob/master/markov chains.ipynb

The drugs/chemicals used in perfusion experiments were:

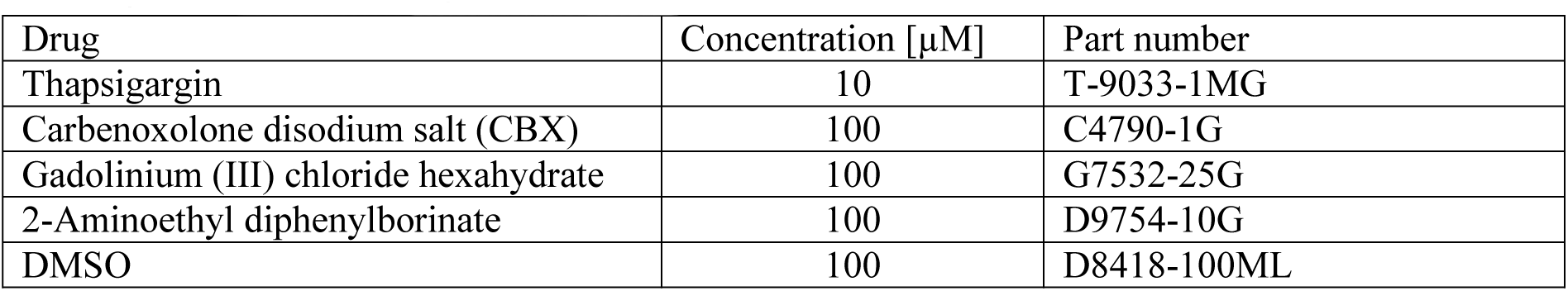

### Drug experiments with cytoskeletal markers

Fertilized eggs were electroporated with one of the following constructs at 80-100μg: Brachyury>mCherry-hsActin, Brachyury> mCherry-Utrophin, Brachyury>mCherry-TalinA, Brachyury> mCherry-MLC, Brachyury>Ensconsin-GFP Transgenics were incubated with control solution (DMSO alone) or drug solution at two different developmental stages. The first set was incubated starting at stage 13 (late gastrula) until they reached stage 24 (late tailbud II). The second set of embryos was incubated from the onset of Stage 24 (late tailbud II) until the end of Stage 25 (late tailbud III). Embryos were fixed at the end of incubation and then imaged using a Leica TSP5 confocal using the following settings: 40x NA1.25 oil objective, HyD detector, 488nm for Brachyury>Ensconsin-GFP or 561nm laser for all other transgenes.

### Quantification of cytoskeletal markers

Stacks acquired with the Leica TSP5 (.lif format) were converted to .tif files. We calculated the fractal dimension of thresholded image stacks using our own 3D-implementation of the box-counting method. The statistical difference between groups was determined with a Mann-Whitney U-test with Holm correction, using the scikit-posthocs python library. For further details on our code and technique, please refer to https://github.com/ChatzigeorgiouGroup/FractalDimension.

### Dm-E-Cadherin marker localization data acquisition and quantification

Fertilized eggs were electroporated with Brachyury>Dm-E-Cadherin-mCherry at 80μg. Embryos were incubated from stage 13 (late gastrula) until they reached stage 24 (late tailbud II) with control solution or drug solutions. They were subsequently fixed and imaged under a Leica TSP5 confocal microscope using a 40x NA1.25 oil objective, a HyD detector and 561nm laser. The collected stacks were the used to generate Maximal projections in ImageJ. We then generated intensity profiles across the lateral and basal membranes demarcated by the Dm-E-cadherin-mCherry expression. Since care was taken during the acquisition of the data to always centre the measurements on the membrane, we could centre the data on peaks by aligning the middle of the intensity profile measurements. We calculated the mean intensity in a 2μm wide band centred on the peak and computed the ratio between the lateral and basal sides mean intensities. Statistical significance in comparisons between groups was determined using Mann-Whitney U tests with Holm correction.

### PACR optogenetic experiments

The expression vector Brachyury>msEGFP-PACR>unc-54 3’UTR was electroporated at 100μg. Electroporated eggs were spread out in 9cm petri dishes in an INCU-Line Tower cooled incubator (maintained at 14°C). This incubator was lined up with 230 Luxorparts Adresserbar RGB WS2812 LED lights (Kjell & Company, part number 87963). For our optogenetic experiments we used the LEDs that delivered 470nm light for illumination. Control plates were wrapped in aluminium foil to prevent blue light reaching the embryos, while embryos that we wanted to be illuminated with the blue light were in optically transparent petri-dishes (with the lids on). Incubation with the 470nm light started after stage 5a (early 16-cell stage) and the embryos were removed and fixed at stage 24 (late tailbud II). Fixed embryos were stained with a conjugated Alexa Fluor 647 phalloidin antibody (1:250). Confocal stacks were collected using a Leica TSP5 confocal with the following conditions: 40x NA1.25 oil objective, PMT detectors and 488nm and 633nm Excitation lasers.

We then generated z-stack maximal projections and using custom made software we labelled notochord cell centres from 1-40 in ascending order along the Anterior-Posterior axis using the msEGFP signal from PACR to identify individual cells. In addition, we segmented the embryo outline using the phalloidin signal in order to calculate the skeleton (the line that defines the body shape of the embryo). For all labelled cells, the distance to both lateral outlines and the central skeleton line was calculated. Only the distance to skeleton was used in the manuscript. Comparisons of the distance to the central skeleton line for each cell number gives an indication of intercalation quality. Mann-Whitney U-tests per cell number were performed to determine statistical difference in distance to the skeleton line for that Dark vs Light groups. For calculating intercellular distances we measured the distance in positions between subsequent notochord cells (cell1→cell2, cell2→3). Then we plotted the mean value of intercellular distance for all embryos.

## Author Contributions

M.C., J.H. and M.S. designed the project and performed the experiments. R.E, K.K, D.D, provided reagents and analysis tools. M.C, J.H, M.S K.K. and D.D, analysed and interpreted data. M.C wrote the manuscript with input from all authors.

## Acknowledgements

We would like to thank members of the Chatzigeorgiou lab and Mie Wong for comments on the manuscript. We are grateful to Dr. Di Jiang for the actin and microtubule markers.

---

**Supplemental Figure 1:**
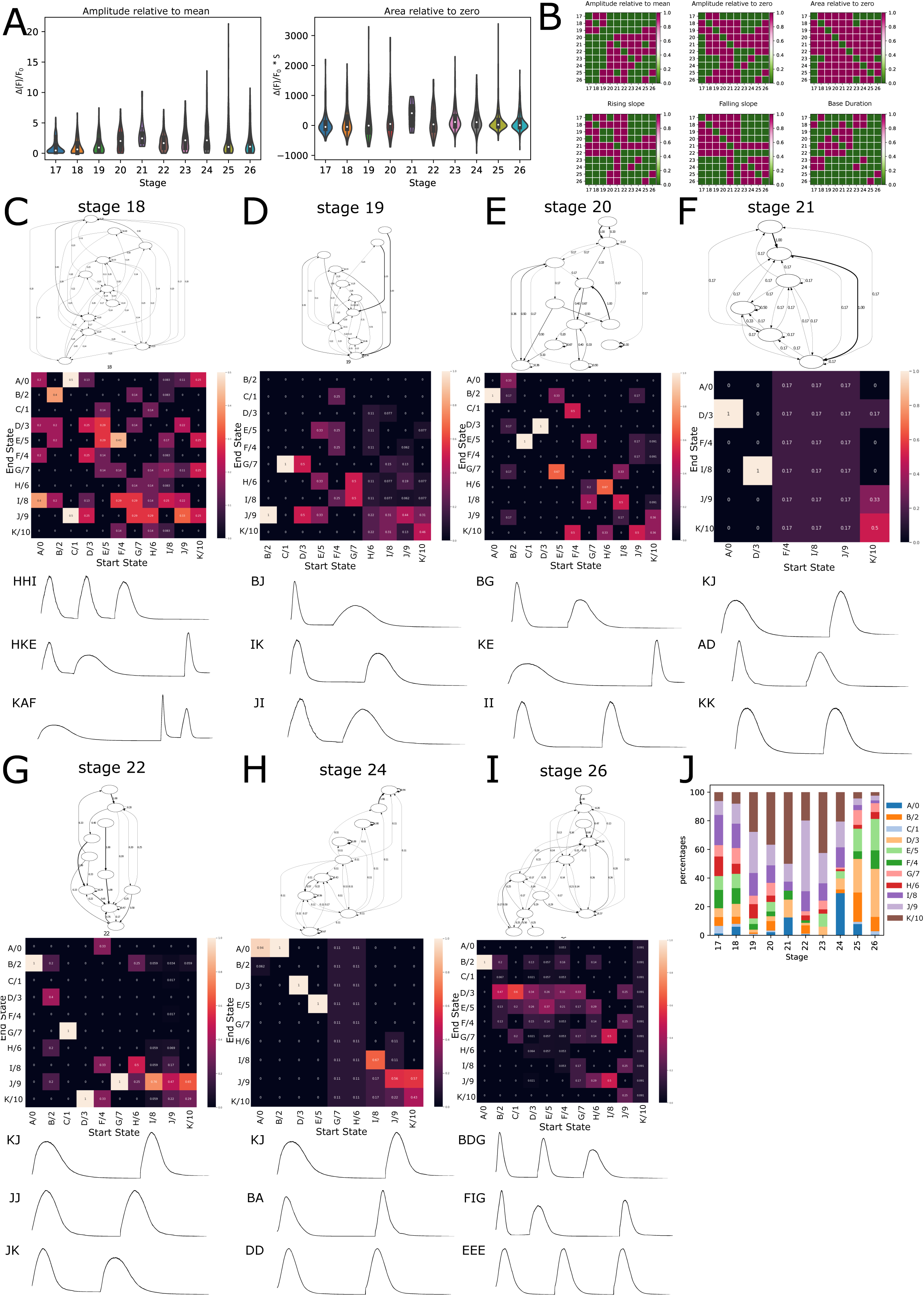
During development notochord cells show a diversity of Ca^2+^ profiles. A) Additional peak features of stage specific Ca^2+^ imaging B) Statistical significance matrices for peak features shown in Figure 1 and Sup. Figure 1. Detailed p-value tables can be found in Sup. Tables 16-21 (C-I) Stage specific Markov-chain generated transition graphs, transition matrices and example traces generated from the Markov Chain model, J) Stage specific k-Shape cluster contribution (%) shown in the form of stacked bar graph. The data shown is the same as in Figure 1F.

**Supplemental Figure 2:**
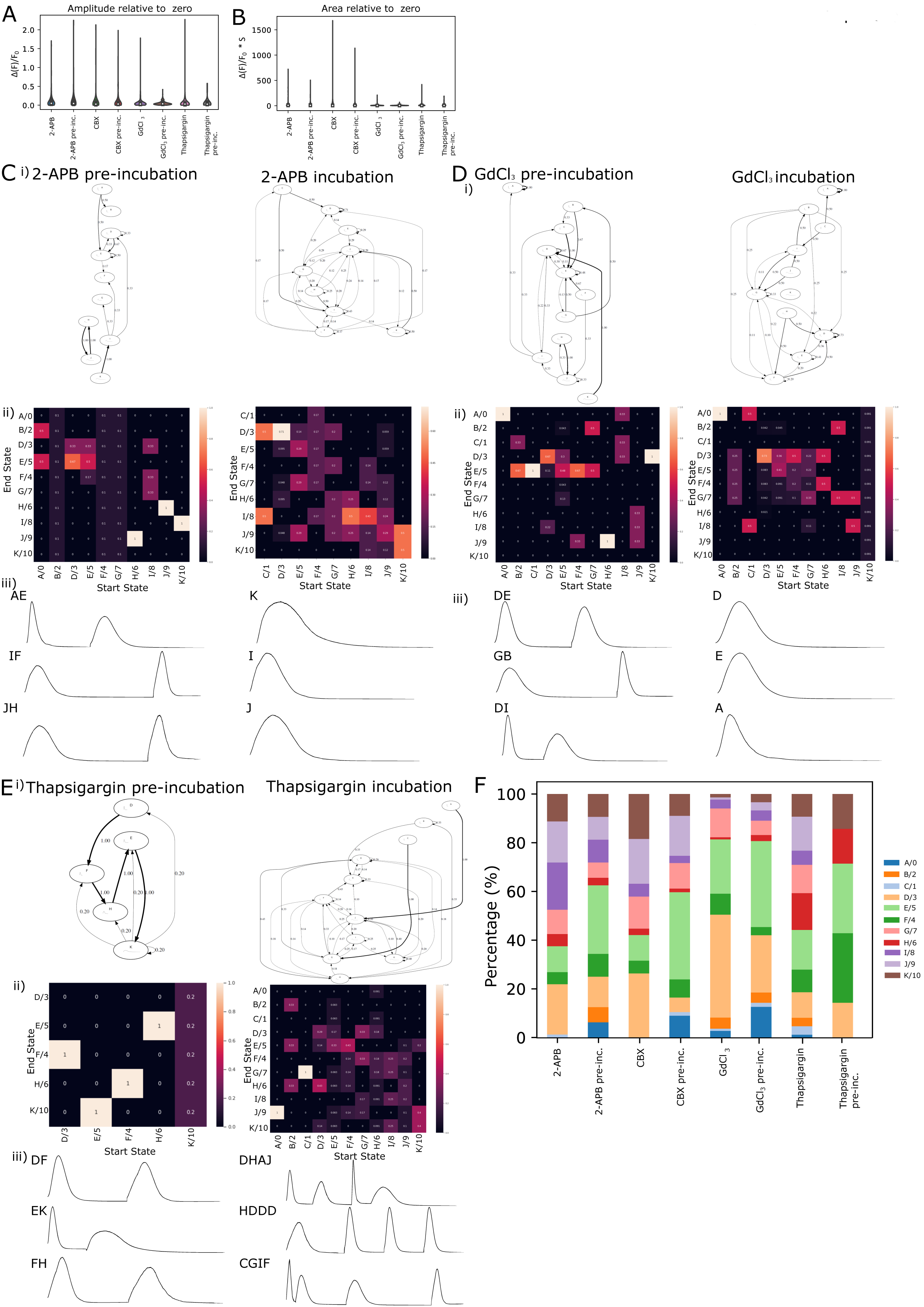
Ca^2+^ signalling dynamics during tubulogenesis are modulated by diverse signalling pathways. (A-B) Additional peak features of Ca^2+^ transients from control and drug treated notochord cells during tubulogenesis: (A) Amplitude relative to zero (B) Area relative to zero. (C-E) (i) Markov-chain generated transition graphs, (ii) transition matrices and (iii) example traces generated from the Markov Chain model, for control and drug treated embryos F) Control and drug treated k-Shape cluster contribution (%) shown in the form of stacked bar graph. The data shown is the same as in Figure 1F.

## Notes

### Competing Interest Statement

The authors have declared no competing interest.

