## Supplemental Information for "Mapping calcium dynamics in a developing tubular structure"

Supplemental Table 1: Calcium imaging dataset information used in Figure 1

| Wild-type dataset | | | |
| --- | --- | --- | --- |
| Stage | Number of animals | Number of curves | UUIDs |
| 17 | 8 | 41 | 205 |
| 18 | 5 | 30 | 96 |
| 19 | 5 | 40 | 101 |
| 20 | 13 | 49 | 89 |
| 21 | 4 | 9 | 16 |
| 22 | 7 | 69 | 172 |
| 23 | 9 | 26 | 186 |
| 24 | 11 | 37 | 168 |
| 25 | 14 | 187 | 679 |
| 26 | 7 | 71 | 261 |

Supplemental Table 2: Calcium imaging dataset information used in Figure 2

| Drugs dataset | | | |
| --- | --- | --- | --- |
| Drug | Number of animals | Number of curves | UUIDs |
| 2-APB (Pre-incubation) | 5 | 11 | 114 |
| 2-APB (Incubation) | 18 | 95 | 470 |
| GdCl3 (Pre-incubation) | 8 | 61 | 303 |
| GdCl3 (Incubation) | 14 | 127 | 761 |
| Thapsigargin (Pre incubation) | 4 | 18 | 28 |
| Thapsigargin (Incubation) | 5 | 5 | 284 |
| CBX (Pre-incubation) | 6 | 33 | 319 |
| CBX (Incubation) | 4 | 27 | 180 |

Supplemental Table 3: Numbers of animals used in Figure 3

| Construct | 2-APB | CBX | Control | GdCl_3_ |
| --- | --- | --- | --- | --- |
| Brachyury>mcherry hs Actin (Gastrula) | 51 | 57 | 63 | 44 |
| Brachyury>mcherry hs Actin (Stg24) | 52 | 63 | 57 | 51 |
| Brachyury>mcherry Utrophin (Gastrula) | 40 | 58 | 38 | 48 |
| Brachyury>mcherry Utrophin (Stg24) | 37 | 38 | 34 | 57 |
| Brachyury>mcherry MLC (Gastrula) | 35 | 13 | 23 | 15 |
| Brachyury>mcherry MLC (Stg24) | 50 | 66 | 61 | 53 |
| Brachyury>Ensconsin GFP (Gastrula) | 70 | 44 | 38 | 17 |
| Brachyury>Ensconsin GFP (Stg24) | 71 | 56 | 25 | 47 |

Supplemental Table 4: p-values for Brachyury>mcherry hs Actin

| Gastrula incubation | 2-APB | CBX | Control | GdCl_3_ |
| --- | --- | --- | --- | --- |
| 2-APB | -1.0 | 0.26305248926221464 | 0.3915821478251902 | 0.26305248926221464 |
| CBX | 0.26305248926221464 | -1.0 | 0.012884707530241162 | 0.9918030846130503 |
| Control | 0.3915821478251902 | 0.012884707530241162 | -1.0 | 0.018120051118599767 |
| GdCl_3_ | 0.26305248926221464 | 0.9918030846130503 | 0.018120051118599767 | -1.0 |
| Stg 24 incubation |  |  |  |  |
| 2-APB | -1.0 | 1.0 | 0.00040862612909187357 | 0.0217264304360778 |
| CBX | 1.0 | -1.0 | 0.00649524646479053 | 0.14984702714672982 |
| Control | 0.00040862612909187357 | 0.00649524646479053 | -1.0 | 1.0 |
| GdCl_3_ | 0.0217264304360778 | 0.14984702714672982 | 1.0 | -1.0 |

Supplemental Table 5: p-values for Brachyury>mcherry Utrophin

| Gastrula incubation | 2-APB | CBX | Control | GdCl3 |
| --- | --- | --- | --- | --- |
| 2-APB | -1.0 | 0.054459363419369834 | 0.18865718666799103 | 1.4326920036056656e-05 |
| CBX | 0.054459363419369834 | -1.0 | 0.009361781357946184 | 1.8753271409810157e-08 |
| Control | 0.18865718666799103 | 0.009361781357946184 | -1.0 | 0.0010936564313249546 |
| GdCl3 | 1.432692e-05 | 1.875327e-08 | 0.0010936564313249546 | -1.0 |
| Stg 24 incubation |  |  |  |  |
| 2-APB | -1.0 | 0.6317346613016089 | 1.4957584588706243e-06 | 0.9530557558838932 |
| CBX | 0.6317346613016089 | -1.0 | 6.281600504012409e-07 | 0.6317346613016089 |
| Control | 1.4957584588706243e-06 | 6.281600504012409e-07 | -1.0 | 8.842396717682183e-07 |
| GdCl_3_ | 0.9530557558838932 | 0.6317346613016089 | 8.842396717682183e-07 | -1.0 |

Supplemental Table 6: p-values for Brachyury>mcherry MLC

| Gastrula incubation | 2-APB | CBX | Control | GdCl_3_ |
| --- | --- | --- | --- | --- |
| 2-APB | -1.0 | 0.10571211879193912 | 1.0 | 0.027339789706911685 |
| CBX | 0.10571211879193912 | -1.0 | 0.10571211879193912 | 1.0 |
| Control | 1.0 | 0.10571211879193912 | -1.0 | 0.9422901219110604 |
| GdCl_3_ | 0.027339789706911685 | 1.0 | 0.10571211879193912 | -1.0 |
| Stg 24 incubation |  |  |  |  |
| 2-APB | -1.0 | 0.007967136854774978 | 4.198457184415616e-05 | 0.0014277777372909463 |
| CBX | 0.007967136854774978 | -1.0 | 8.382251416550418e-11 | 2.2768062178794417e-08 |
| Control | 4.198457184415616e-05 | 8.382251416550418e-11 | -1.0 | 0.4397348636077315 |
| GdCl_3_ | 0.0014277777372909 | 2.2768062178794417e-08 | 0.4397348636077315 | -1.0 |

Supplemental Table 7: p-values for Brachyury>Ensconsin GFP

| Gastrula incubation | 2-APB | CBX | Control | GdCl_3_ |
| --- | --- | --- | --- | --- |
| 2-APB | -1.0 | 1.0 | 1.0 | 0.00017424157319935333 |
| CBX | 1.0 | -1.0 | 1.0 | 0.0031513422013594024 |
| Control | 1.0 | 1.0 | -1.0 | 0.030503764947027744 |
| GdCl_3_ | 0.00017424157319935333 | 0.0031513422013594024 | 0.030503764947027744 | -1.0 |
| Stg 24 incubation |  |  |  |  |
| 2-APB | -1.0 | 0.30699500692269255 | 0.30699500692269255 | 0.4701489320559986 |
| CBX | 0.30699500692269255 | -1.0 | 0.05238020204866552 | 0.6312055601669118 |
| Control | 0.30699500692269255 | 0.05238020204866552 | -1.0 | 0.13903353518988426 |
| GdCl_3_ | 0.4701489320559986 | 0.6312055601669118 | 0.13903353518988426 | -1.0 |

Supplemental Table 8: Number of measurements DeActs

| DeActs Experiment | | | |
| --- | --- | --- | --- |
| Condition | Number of animals | Number of curves | UUIDs |
| Control | 24 | 74 | 233 |
| DeActs | 32 | 116 | 964 |

Supplemental Table 9: Number of measurements Dm-E-Cadherin>mcherry

|  | Lateral | Basal |
| --- | --- | --- |
| 2-APB | 69 | 68 |
| CBX | 61 | 61 |
| Control | 69 | 68 |
| GdCl_3_ | 40 | 41 |

Supplemental Table 10: p-values for Dm-E-Cadherin>mcherry lateral versus basal intensity ratio

|  | GdCl_3_ | CBX | Control | 2-APB |
| --- | --- | --- | --- | --- |
| GdCl_3_ | -1.0 | 0.23686932912938313 | 0.0036951708378875456 | 0.9798807385120032 |
| CBX | 0.23686932912938313 | -1.0 | 0.022810833980056212 | 0.23686932912938313 |
| Control | 0.0036951708378875456 | 0.022810833980056212 | -1.0 | 0.00040623001992318576 |
| 2-APB | 0.9798807385120032 | 0.23686932912938313 | 0.00040623001992318576 | -1.0 |

Supplemental Table11: Forward vs Reverse Bias for Figure 1 and Sup. Figure 1 data

| Wild-type dataset | | |
| --- | --- | --- |
| Condition | Forward Bias | Reverse Bias |
| Stage 17 | 5.30 | 4.46 |
| Stage 18 | 4.75 | 4.39 |
| Stage 19 | 6.21 | 2.12 |
| Stage 20 | 5.27 | 3.04 |
| Stage 21 | 3.00 | 2.00 |
| Stage 22 | 7.26 | 1.72 |
| Stage 23 | 3.19 | 1.92 |
| Stage 24 | 1.40 | 2.79 |
| Stage 25 | 2.06 | 5.16 |
| Stage 26 | 5.09 | 4.68 |

Supplemental Table12: Forward vs Reverse Bias for Figure 2 and Sup. Figure 2 data

| Drugs | | |
| --- | --- | --- |
| Condition | Forward Bias | Reverse Bias |
| 2-APB (Pre-incubation) | 4.53 | 4.33 |
| 2-APB (Incubation) | 3.94 | 2.42 |
| CBX (Pre-incubation) | 4.11 | 4.06 |
| CBX (Incubation) | 2.68 | 3.62 |
| GdCl3 (Pre-incubation) | 3.84 | 4.68 |
| GdCl3 (Incubation) | 2.12 | 6.12 |
| Thapsigargin (Pre incubation) | 3.00 | 1.80 |
| Thapsigargin (Incubation) | 5.01 | 4.06 |

Supplemental Table13: Forward vs Reverse Bias for Figure 4 data

| DeActs Experiment | | |
| --- | --- | --- |
| Condition | Forward Bias | Reverse Bias |
| Control | 4.42 | 2.42 |
| Cah3>DeAct-SpvB | 5.88 | 3.77 |

Supplemental Table14: p-values for Peak features for Figure 4 data

| Feature | Amplitude relative to zero | Area relative to zero | Rising slope | Falling slope | Base duration |
| --- | --- | --- | --- | --- | --- |
| p-value | 1.4427547353400655e-19 | 2.5041038359344278e-26 | 6.623532982712519e-11 | 6.320865097759602e-13 | 3.363230739648894e-08 |

Supplemental Table 15: p-values for Peak features in Figure 2 and Sup. Figure 2

| Feature | Amplitude relative to mean | Amplitude relative to zero | Area relative to zero | Rising slope | Falling slope | Base duration |
| --- | --- | --- | --- | --- | --- | --- |
| Thapsigargin | 0.4058484714885624 | 0.3354638785536568 | 0.2253538913106511 | 0.3110264676076546 | 0.40159680543512305 | 0.49956200489143554 |
| 2-APB | 0.30670747668707543 | 0.356264262094361 | 0.482603062472704 | 0.24619987862304998 | 0.3727678472963607 | 0.08661797999503124 |
| GdCl3 | 0.007000109786320786 | 0.001438904575296533 | 0.00022938949770633115 | 0.008157921109204698 | 0.013617685603329209 | 0.46520916838442844 |
| CBX | 0.05737973502960664 | 0.035508080371025194 | 0.002945373225743248 | 0.23799241458443898 | 0.184506654652694 | 2.802032429238443e-06 |

Supplemental Table 16: P-values for Wild-type dataset Rising Slope (Slope of the line drawn from the left base to the peak)

|  | 17 | 18 | 19 | 20 | 21 | 22 | 23 | 24 | 25 | 26 |
| --- | --- | --- | --- | --- | --- | --- | --- | --- | --- | --- |
| 17 | -1.0 | 1.0 | 1.0 | 0.012492829850752629 | 0.2670940654707191 | 0.07392550993823369 | 9.835040972309486e-25 | 5.889041478681398e-17 | 4.690558674701061e-19 | 3.243178857061207e-11 |
| 18 | 1.0 | -1.0 | 1.0 | 0.0864235667852451 | 0.2670940654707191 | 0.7939571739711885 | 2.9775294916754834e-13 | 7.799656520389342e-12 | 1.0287909352194868e-09 | 5.703929836583393e-07 |
| 19 | 1.0 | 1.0 | -1.0 | 0.016373599316802143 | 0.2670940654707191 | 0.13427341777589172 | 6.51437343831652e-18 | 5.121822763129666e-13 | 3.2232257771059265e-12 | 3.059517563461333e-08 |
| 20 | 0.012492829850752629 | 0.0864235667852451 | 0.016373599316802143 | -1.0 | 1.0 | 1.0 | 0.00011062449360080593 | 0.0019597151988287654 | 0.18345946319532924 | 0.7845259210837155 |
| 21 | 0.2670940654707191 | 0.2670940654707191 | 0.2670940654707191 | 1.0 | -1.0 | 1.0 | 1.0 | 1.0 | 1.0 | 1.0 |
| 22 | 0.07392550993823369 | 0.7939571739711885 | 0.13427341777589172 | 1.0 | 1.0 | -1.0 | 2.270627054882447e-17 | 6.50170536512239e-10 | 5.650750333552378e-09 | 8.212305618379863e-05 |
| 23 | 9.835040972309486e-25 | 2.9775294916754834e-13 | 6.51437343831652e-18 | 0.00011062449360080593 | 1.0 | 2.270627054882447e-17 | -1.0 | 1.0 | 0.01282143630414814 | 0.019837915186544534 |
| 24 | 5.889041478681398e-17 | 7.799656520389342e-12 | 5.121822763129666e-13 | 0.0019597151988287654 | 1.0 | 6.50170536512239e-10 | 1.0 | -1.0 | 0.026540454717360433 | 0.04026041579328745 |
| 25 | 4.690558674701061e-19 | 1.0287909352194868e-09 | 3.2232257771059265e-12 | 0.18345946319532924 | 1.0 | 5.650750333552378e-09 | 0.01282143630414814 | 0.026540454717360433 | -1.0 | 1.0 |
| 26 | 3.243178857061207e-11 | 5.703929836583393e-07 | 3.059517563461333e-08 | 0.7845259210837155 | 1.0 | 8.212305618379863e-05 | 0.019837915186544534 | 0.04026041579328745 | 1.0 | -1.0 |

Supplemental Table 17: P-values for Wild-type dataset Falling Slope

|  | 17 | 18 | 19 | 20 | 21 | 22 | 23 | 24 | 25 | 26 |
| --- | --- | --- | --- | --- | --- | --- | --- | --- | --- | --- |
| 17 | -1.0 | 1.0 | 1.0 | 1.0 | 0.24389344096362917 | 0.3964055977886872 | 4.399940791334594e-14 | 2.5872138514678975e-09 | 2.300481469365248e-09 | 1.4916081704174834e-07 |
| 18 | 1.0 | -1.0 | 1.0 | 0.9804184814628945 | 0.5454438549958879 | 0.7165175807248667 | 1.659392651479232e-08 | 1.5635695356728143e-07 | 5.66833325545375e-07 | 1.1131696873669104e-05 |
| 19 | 1.0 | 1.0 | -1.0 | 0.7165175807248667 | 0.15435531611038864 | 0.06642264135347906 | 2.172882355189465e-12 | 3.899623531007742e-09 | 1.375938939757955e-08 | 2.1754099663410955e-07 |
| 20 | 1.0 | 0.9804184814628945 | 0.7165175807248667 | -1.0 | 1.0 | 1.0 | 2.823840886498772e-05 | 0.00169744534772009 | 0.02712155282987664 | 0.025896400986432436 |
| 21 | 0.24389344096362917 | 0.5454438549958879 | 0.15435531611038864 | 1.0 | -1.0 | 1.0 | 1.0 | 1.0 | 1.0 | 1.0 |
| 22 | 0.3964055977886872 | 0.7165175807248667 | 0.06642264135347906 | 1.0 | 1.0 | -1.0 | 6.370715867449619e-09 | 5.030238024853173e-05 | 0.0015987468291324793 | 0.0021287627800175373 |
| 23 | 4.399940791334594e-14 | 1.659392651479232e-08 | 2.172882355189465e-12 | 2.823840886498772e-05 | 1.0 | 6.370715867449619e-09 | -1.0 | 1.0 | 0.2599645063959843 | 0.9804184814628945 |
| 24 | 2.5872138514678975e-09 | 1.5635695356728143e-07 | 3.899623531007742e-09 | 0.00169744534772009 | 1.0 | 5.030238024853173e-05 | 1.0 | -1.0 | 0.7845036686700535 | 1.0 |
| 25 | 2.300481469365248e-09 | 5.66833325545375e-07 | 1.375938939757955e-08 | 0.02712155282987664 | 1.0 | 0.0015987468291324793 | 0.2599645063959843 | 0.7845036686700535 | -1.0 | 1.0 |
| 26 | 1.4916081704174834e-07 | 1.1131696873669104e-05 | 2.1754099663410955e-07 | 0.025896400986432436 | 1.0 | 0.0021287627800175373 | 0.9804184814628945 | 1.0 | 1.0 | -1.0 |

Supplemental Table 18: P-values for Wild-type dataset Base Duration

|  | 17 | 18 | 19 | 20 | 21 | 22 | 23 | 24 | 25 | 26 |
| --- | --- | --- | --- | --- | --- | --- | --- | --- | --- | --- |
| 17 | -1.0 | 1.0 | 5.789875148295449e-13 | 3.600353887125212e-10 | 0.006596544716436981 | 5.225866373532579e-22 | 1.0 | 1.0 | 0.0015621241690529637 | 0.002949614851125152 |
| 18 | 1.0 | -1.0 | 1.316484674612523e-06 | 1.934322817978465e-05 | 0.05192318636451625 | 6.058768500009934e-11 | 1.0 | 1.0 | 0.003135868229522879 | 0.003397153755470576 |
| 19 | 5.789875148295449e-13 | 1.316484674612523e-06 | -1.0 | 1.0 | 1.0 | 1.0 | 3.549277211307667e-12 | 6.802168500384217e-10 | 1.9495585490199737e-26 | 4.503912040513792e-25 |
| 20 | 3.600353887125212e-10 | 1.934322817978465e-05 | 1.0 | -1.0 | 1.0 | 1.0 | 5.524418827828403e-09 | 1.760894214409854e-07 | 3.637251528595568e-20 | 4.914303203079985e-19 |
| 21 | 0.006596544716436981 | 0.05192318636451625 | 1.0 | 1.0 | -1.0 | 1.0 | 0.017671846144453364 | 0.019386946113637425 | 0.0010495583786702544 | 0.000800665901084004 |
| 22 | 5.225866373532579e-22 | 6.058768500009934e-11 | 1.0 | 1.0 | 1.0 | -1.0 | 4.143931743351004e-20 | 1.5435555196178076e-16 | 7.457031493941256e-48 | 3.942542281013977e-41 |
| 23 | 1.0 | 1.0 | 3.549277211307667e-12 | 5.524418827828403e-09 | 0.017671846144453364 | 4.143931743351004e-20 | -1.0 | 1.0 | 1.9505059216990236e-07 | 2.4843907547733945e-06 |
| 24 | 1.0 | 1.0 | 6.802168500384217e-10 | 1.760894214409854e-07 | 0.019386946113637425 | 1.5435555196178076e-16 | 1.0 | -1.0 | 2.3408763398598062e-09 | 1.3193012829654109e-08 |
| 25 | 0.0015621241690529637 | 0.003135868229522879 | 1.9495585490199737e-26 | 3.637251528595568e-20 | 0.0010495583786702544 | 7.457031493941256e-48 | 1.9505059216990236e-07 | 2.3408763398598062e-09 | -1.0 | 1.0 |
| 26 | 0.002949614851125152 | 0.003397153755470576 | 4.503912040513792e-25 | 4.914303203079985e-19 | 0.000800665901084004 | 3.942542281013977e-41 | 2.4843907547733945e-06 | 1.3193012829654109e-08 | 1.0 | -1.0 |

Supplemental Table 19: P-values area under the curve relative to zero

|  | 17 | 18 | 19 | 20 | 21 | 22 | 23 | 24 | 25 | 26 |
| --- | --- | --- | --- | --- | --- | --- | --- | --- | --- | --- |
| 17 | -1.0 | 1.0 | 1.0 | 0.15997287651091077 | 0.16286121759928102 | 1.0 | 4.42337155404124e-08 | 1.273413576507818e-06 | 8.698913452113917e-08 | 0.0017211613423960051 |
| 18 | 1.0 | -1.0 | 1.0 | 1.0 | 1.0 | 1.0 | 0.04289860250215418 | 0.07009996297395298 | 0.11921291757850704 | 1.0 |
| 19 | 1.0 | 1.0 | -1.0 | 1.0 | 1.0 | 1.0 | 0.1678024894197146 | 0.3233999116110465 | 0.9081865537566806 | 1.0 |
| 20 | 0.15997287651091077 | 1.0 | 1.0 | -1.0 | 1.0 | 1.0 | 1.0 | 1.0 | 1.0 | 1.0 |
| 21 | 0.16286121759928102 | 1.0 | 1.0 | 1.0 | -1.0 | 1.0 | 1.0 | 1.0 | 0.9081865537566806 | 0.8804937973314163 |
| 22 | 1.0 | 1.0 | 1.0 | 1.0 | 1.0 | -1.0 | 0.8601913679827828 | 1.0 | 1.0 | 1.0 |
| 23 | 4.42337155404124e-08 | 0.04289860250215418 | 0.1678024894197146 | 1.0 | 1.0 | 0.8601913679827828 | -1.0 | 1.0 | 0.05435459416764868 | 0.037805228945025464 |
| 24 | 1.273413576507818e-06 | 0.07009996297395298 | 0.3233999116110465 | 1.0 | 1.0 | 1.0 | 1.0 | -1.0 | 0.10993591478121365 | 0.07070102582764333 |
| 25 | 8.698913452113917e-08 | 0.11921291757850704 | 0.9081865537566806 | 1.0 | 0.9081865537566806 | 1.0 | 0.05435459416764868 | 0.10993591478121365 | -1.0 | 1.0 |
| 26 | 0.0017211613423960051 | 1.0 | 1.0 | 1.0 | 0.8804937973314163 | 1.0 | 0.037805228945025464 | 0.07070102582764333 | 1.0 | -1.0 |

Supplemental Table 20: P-value Peak Amplitude relative to zero

|  | 17 | 18 | 19 | 20 | 21 | 22 | 23 | 24 | 25 | 26 |
| --- | --- | --- | --- | --- | --- | --- | --- | --- | --- | --- |
| 17 | -1.0 | 1.0 | 0.8967923041465762 | 3.4864161591662277e-06 | 0.0007504243580555914 | 0.00046012418794200936 | 5.360219906200857e-16 | 6.4196959238441426e-15 | 4.272403039866185e-11 | 2.3913907090179208e-05 |
| 18 | 1.0 | -1.0 | 1.0 | 0.06612184156860354 | 0.06700371381194395 | 1.0 | 1.262698866612348e-05 | 6.746182637128074e-06 | 0.014072765234424597 | 0.541141973553521 |
| 19 | 0.8967923041465762 | 1.0 | -1.0 | 0.09251925995031242 | 0.06855046154450732 | 1.0 | 4.7462827471719116e-05 | 1.3213199962574223e-05 | 0.06855046154450732 | 0.729878724446098 |
| 20 | 3.4864161591662277e-06 | 0.06612184156860354 | 0.09251925995031242 | -1.0 | 1.0 | 0.5709934013517766 | 1.0 | 0.5139038486132126 | 1.0 | 1.0 |
| 21 | 0.0007504243580555914 | 0.06700371381194395 | 0.06855046154450732 | 1.0 | -1.0 | 0.11485508713790726 | 1.0 | 1.0 | 0.20186793942678713 | 0.20314238882858285 |
| 22 | 0.00046012418794200936 | 1.0 | 1.0 | 0.5709934013517766 | 0.11485508713790726 | -1.0 | 0.0013372185081187858 | 0.0002845698467662035 | 1.0 | 1.0 |
| 23 | 5.360219906200857e-16 | 1.262698866612348e-05 | 4.7462827471719116e-05 | 1.0 | 1.0 | 0.0013372185081187858 | -1.0 | 1.0 | 0.0014321972567762737 | 0.0039426171524547015 |
| 24 | 6.4196959238441426e-15 | 6.746182637128074e-06 | 1.3213199962574223e-05 | 0.5139038486132126 | 1.0 | 0.0002845698467662035 | 1.0 | -1.0 | 0.0005510795772384252 | 0.002176688062302675 |
| 25 | 4.272403039866185e-11 | 0.014072765234424597 | 0.06855046154450732 | 1.0 | 0.20186793942678713 | 1.0 | 0.0014321972567762737 | 0.0005510795772384252 | -1.0 | 1.0 |
| 26 | 2.3913907090179208e-05 | 0.541141973553521 | 0.729878724446098 | 1.0 | 0.20314238882858285 | 1.0 | 0.0039426171524547015 | 0.002176688062302675 | 1.0 | -1.0 |

Supplemental Table 21: P-values for Peak Amplitude relative to the mean of its bases

|  | 17 | 18 | 19 | 20 | 21 | 22 | 23 | 24 | 25 | 26 |
| --- | --- | --- | --- | --- | --- | --- | --- | --- | --- | --- |
| 17 | -1.0 | 1.0 | 0.2550900966214597 | 3.356113478170455e-07 | 3.195559153535446e-05 | 1.5525987158947807e-12 | 7.486653029947438e-20 | 2.4296736502674033e-16 | 5.059652976448185e-09 | 2.9344468738973818e-06 |
| 18 | 1.0 | -1.0 | 0.2550900966214597 | 1.3421851532165434e-05 | 0.000335700321907161 | 1.3352626598252904e-06 | 8.526645436977498e-11 | 2.97584648359941e-10 | 3.8056636870756825e-05 | 0.0004445562222751226 |
| 19 | 0.2550900966214597 | 0.2550900966214597 | -1.0 | 0.026870840029626884 | 0.005049288954035554 | 0.011698806158533146 | 5.641137304646892e-06 | 1.1818403016223825e-05 | 0.5115832680923265 | 0.7407555385446153 |
| 20 | 3.356113478170455e-07 | 1.3421851532165434e-05 | 0.026870840029626884 | -1.0 | 0.8759574001272723 | 1.0 | 1.0 | 0.7796517743817206 | 0.4630730033733127 | 0.5144588153118801 |
| 21 | 3.195559153535446e-05 | 0.000335700321907161 | 0.005049288954035554 | 0.8759574001272723 | -1.0 | 0.16270547160506035 | 1.0 | 1.0 | 0.045423059211086914 | 0.05242807356671181 |
| 22 | 1.5525987158947807e-12 | 1.3352626598252904e-06 | 0.011698806158533146 | 1.0 | 0.16270547160506035 | -1.0 | 0.2550900966214597 | 0.15946346471281672 | 0.1319334980089507 | 0.2550900966214597 |
| 23 | 7.486653029947438e-20 | 8.526645436977498e-11 | 5.641137304646892e-06 | 1.0 | 1.0 | 0.2550900966214597 | -1.0 | 1.0 | 6.179462715058664e-06 | 0.00021473248945253246 |
| 24 | 2.4296736502674033e-16 | 2.97584648359941e-10 | 1.1818403016223825e-05 | 0.7796517743817206 | 1.0 | 0.15946346471281672 | 1.0 | -1.0 | 2.9993628989278182e-05 | 0.0004408069351353227 |
| 25 | 5.059652976448185e-09 | 3.8056636870756825e-05 | 0.5115832680923265 | 0.4630730033733127 | 0.045423059211086914 | 0.1319334980089507 | 6.179462715058664e-06 | 2.9993628989278182e-05 | -1.0 | 1.0 |
| 26 | 2.9344468738973818e-06 | 0.0004445562222751226 | 0.7407555385446153 | 0.5144588153118801 | 0.05242807356671181 | 0.2550900966214597 | 0.00021473248945253246 | 0.0004408069351353227 | 1.0 | -1.0 |

Supplemental Table 22: Mann-Whitney values per cell number for Figure 2D

|  | cell_number | p_dts | p_dtc | p_dts_bonf | p_dtc_bonf | p_dts_bonf_sig | p_dtc_bonf_sig |
| --- | --- | --- | --- | --- | --- | --- | --- |
| 0 | 1 | 0.443627 | 0.005532 | 17.74509 | 0.221286 | FALSE | FALSE |
| 1 | 2 | 0.162471 | 0.02718 | 6.498856 | 1.087182 | FALSE | FALSE |
| 2 | 3 | 0.000969 | 0.0012 | 0.038743 | 0.047992 | TRUE | TRUE |
| 3 | 4 | 0.000167 | 0.000324 | 0.006661 | 0.012948 | TRUE | TRUE |
| 4 | 5 | 0.000118 | 0.000247 | 0.00471 | 0.00989 | TRUE | TRUE |
| 5 | 6 | 9.86E-06 | 5.73E-05 | 0.000394 | 0.00229 | TRUE | TRUE |
| 6 | 7 | 4.64E-05 | 0.000235 | 0.001855 | 0.009387 | TRUE | TRUE |
| 7 | 8 | 1.00E-05 | 4.44E-05 | 0.0004 | 0.001776 | TRUE | TRUE |
| 8 | 9 | 7.48E-07 | 1.27E-06 | 2.99E-05 | 5.10E-05 | TRUE | TRUE |
| 9 | 10 | 2.52E-09 | 8.62E-09 | 1.01E-07 | 3.45E-07 | TRUE | TRUE |
| 10 | 11 | 6.90E-05 | 0.000236 | 0.002759 | 0.00942 | TRUE | TRUE |
| 11 | 12 | 5.75E-06 | 1.65E-05 | 0.00023 | 0.00066 | TRUE | TRUE |
| 12 | 13 | 4.43E-06 | 1.54E-05 | 0.000177 | 0.000616 | TRUE | TRUE |
| 13 | 14 | 5.89E-09 | 1.84E-08 | 2.35E-07 | 7.36E-07 | TRUE | TRUE |
| 14 | 15 | 1.76E-07 | 6.35E-07 | 7.02E-06 | 2.54E-05 | TRUE | TRUE |
| 15 | 16 | 1.37E-06 | 5.94E-06 | 5.50E-05 | 0.000238 | TRUE | TRUE |
| 16 | 17 | 2.40E-06 | 4.86E-06 | 9.59E-05 | 0.000194 | TRUE | TRUE |
| 17 | 18 | 8.45E-06 | 1.86E-05 | 0.000338 | 0.000744 | TRUE | TRUE |
| 18 | 19 | 5.43E-06 | 2.02E-05 | 0.000217 | 0.000807 | TRUE | TRUE |
| 19 | 20 | 9.65E-06 | 2.50E-05 | 0.000386 | 0.001 | TRUE | TRUE |
| 20 | 21 | 5.12E-07 | 1.13E-06 | 2.05E-05 | 4.52E-05 | TRUE | TRUE |
| 21 | 22 | 3.61E-08 | 1.12E-07 | 1.44E-06 | 4.50E-06 | TRUE | TRUE |
| 22 | 23 | 5.88E-08 | 5.35E-07 | 2.35E-06 | 2.14E-05 | TRUE | TRUE |
| 23 | 24 | 1.09E-06 | 1.16E-05 | 4.36E-05 | 0.000463 | TRUE | TRUE |
| 24 | 25 | 1.95E-06 | 6.51E-06 | 7.81E-05 | 0.00026 | TRUE | TRUE |
| 25 | 26 | 1.91E-08 | 1.65E-07 | 7.63E-07 | 6.60E-06 | TRUE | TRUE |
| 26 | 27 | 2.51E-06 | 1.56E-05 | 0.0001 | 0.000624 | TRUE | TRUE |
| 27 | 28 | 3.52E-06 | 1.52E-05 | 0.000141 | 0.000606 | TRUE | TRUE |
| 28 | 29 | 2.82E-08 | 1.03E-07 | 1.13E-06 | 4.10E-06 | TRUE | TRUE |
| 29 | 30 | 8.16E-08 | 1.81E-06 | 3.26E-06 | 7.24E-05 | TRUE | TRUE |
| 30 | 31 | 6.89E-06 | 5.60E-05 | 0.000276 | 0.002239 | TRUE | TRUE |
| 31 | 32 | 4.37E-07 | 8.08E-06 | 1.75E-05 | 0.000323 | TRUE | TRUE |
| 32 | 33 | 2.53E-09 | 7.93E-08 | 1.01E-07 | 3.17E-06 | TRUE | TRUE |
| 33 | 34 | 2.02E-09 | 1.79E-07 | 8.10E-08 | 7.18E-06 | TRUE | TRUE |
| 34 | 35 | 2.60E-08 | 2.65E-07 | 1.04E-06 | 1.06E-05 | TRUE | TRUE |
| 35 | 36 | 6.17E-12 | 1.25E-10 | 2.47E-10 | 5.00E-09 | TRUE | TRUE |
| 36 | 37 | 2.70E-08 | 1.39E-07 | 1.08E-06 | 5.56E-06 | TRUE | TRUE |
| 37 | 38 | 6.71E-08 | 3.05E-07 | 2.69E-06 | 1.22E-05 | TRUE | TRUE |
| 38 | 39 | 5.78E-07 | 3.65E-07 | 2.31E-05 | 1.46E-05 | TRUE | TRUE |
| 39 | 40 | 6.87E-05 | 1.75E-05 | 0.002747 | 0.000702 | TRUE | TRUE |
